# M.FcoHsdM-dependent DNA methylation coordinates secretion-associated gene expression, interbacterial competition, and virulence in *Flavobacterium columnare*

**DOI:** 10.64898/2026.09.12.750872

**Authors:** Ruoxi Zhu, Yuxuan Zhang, Yongtao Zhu, Runsheng Li, Wenlong Cai

## Abstract

*Flavobacterium columnare* causes columnaris disease in freshwater fish, but the contribution of DNA methylation to its physiology and pathogenicity remains poorly understood. Here, we characterized M. FcoHsdM, a type I restriction-modification methyltransferase previously identified by nanopore methylome analysis. Deletion of the gene encoding M. FcoHsdM abolished methylation at its associated recognition motifs without affecting planktonic growth. The mutant exhibited impaired biofilm formation, gliding motility, colony spreading, and gelatinolytic activity, together with increased outer-membrane vesicle production and complete attenuation in a zebrafish infection model. Transcriptomic and extracellular proteomic analyses revealed broad changes in secretion and cell surface associated functions. In particular, genes encoding components of a putative type VI secretion system were coordinately downregulated, and the mutant showed reduced antagonistic activity against *Aeromonas hydrophila*. Several predicted type IX secretion system substrates and extracellular enzymes were also reduced, potentially contributing to the motility and virulence defects. Two such motifs were identified in *tssP*, which encodes a predicted Bacteroidota-specific type VI secretion system membrane-complex component. One motif overlapped a computationally predicted binding site for the leucine-responsive regulatory protein Lrp, raising the hypothesis that intragenic methylation may influence regulator occupancy at the *tssP* locus. Collectively, these findings identify M.FcoHsdM-dependent methylation as an important contributor to secretion-associated gene expression, interbacterial antagonism, and virulence in *F. columnare*.

## INTRODUCTION

*Flavobacterium columnare* is an opportunistic bacterial pathogen that poses a serious threat to freshwater aquaculture systems worldwide. The pathogen colonizes external mucosal surfaces, especially the gills, skin, and fins, and infection can result in gill necrosis, skin lesions, fin erosion, and mortality [1, 2]. Although host susceptibility and environmental risk factors have been extensively studied, the bacterial regulatory mechanisms that coordinate surface colonization, ecological fitness, and virulence remain incompletely understood.

Protein secretion is central to *F. columnare* pathogenesis. The type IX secretion system (T9SS) characteristic of Bacteroidota exports critical extracellular and surface proteins, including adhesins, enzymes, and gliding motility determinants [3]. Genetic studies have shown that deleting the core T9SS gene *gldN* abolishes secretion, motility, and virulence, and that disrupting *porV* to restrict export of specific T9SS substrates yields comparable attenuation [4–6]. Beyond host infection, *F. columnare* must survive in polymicrobial aquatic environments. The type VI secretion system (T6SS) is a contractile nanomachine that delivers toxic effectors into neighbouring bacterial cells and mediates interbacterial competition [7]. Bacteroidota T6SS loci possess genetic architectures distinct from canonical proteobacterial systems, and recent structural work identified TssNQOPR as a Bacteroidota-specific membrane complex required for T6SS function [8, 9]. Although putative T6SS loci can be identified in some *F. columnare* genomes, their contribution to interbacterial competition and regulatory control remains poorly characterized.

Epigenetic modification via DNA methylation provides an additional regulatory layer governing bacterial physiology. The primary bacterial methylation marks comprise N6-methyladenine (m6A), N4-methylcytosine (m4C), and 5-methylcytosine (5mC) [10, 11]. Although classically viewed as components of restriction-modification systems safeguarding host DNA, bacterial methyltransferases also modulate critical processes including replication, DNA repair, transcription, phase variation, and virulence [12–16]. In type I restriction-modification systems, the HsdS subunit dictates DNA recognition specificity and the HsdM subunit catalyses methyl transfer [17]. Type I methylation patterns can function as regulatory signals that alter DNA-protein interactions [18–21]. A classic example is Dam-dependent methylation at the *Escherichia coli* pap regulatory region, which modulates binding of the leucine-responsive regulatory protein Lrp and generates phase-variable expression [22, 23].

Despite these advances, epigenetic regulation in *F. columnare* has not been investigated. A previous nanopore methylome analysis of *F. columnare* HLCZX-1 identified M. FcoHsdM as a candidate type I methyltransferase. However, its physiological and regulatory functions were unknown. Here, we generated an *M. FcoHsdM* deletion mutant and used integrated multi-omics profiling to determine how M. FcoHsdM-dependent methylation controls *F. columnare* biology. The results show that M.FcoHsdM is required for multiple surface and secretion-associated phenotypes, virulence in zebrafish, and efficient antagonism against *Aeromonas hydrophila*. A putative T6SS locus was coordinately repressed in the mutant, and M.FcoHsdM-dependent methylation sites were identified within the coding sequence of *tssP*. These findings establish M.FcoHsdM as a regulator of pathogenicity-associated traits and identify a candidate connection between intragenic methylation and T6SS-associated gene expression.

## RESULTS

### M. FcoHsdM is responsible for a major methylation signature in *F. columnare*

To investigate the biological role of M. FcoHsdM, an in-frame deletion mutant of the gene encoding this methyltransferase was generated by allelic exchange [24]. Upstream (1880 bp) and downstream (1301 bp) flanking regions of *M. FcoHsdM* were amplified and assembled into the *sacB*-containing suicide vector pMS75, generating plasmid pRX03 (Fig. 1A and Table S3). Following conjugation into the wild-type (WT) strain, tetracycline-resistant integrants were counterselected on sucrose, and recombinants were screened by PCR to confirm deletion of the target gene (Fig. 1B). A complemented strain (C-M. FcoHsdM) was generated by introducing a 2361-bp fragment containing *M. FcoHsdM* and its native promoter into the shuttle vector pCP23.

To determine whether deletion of *M. FcoHsdM* eliminated the corresponding methyltransferase activity, we profiled methylation by nanopore sequencing. The M. FcoHsdM-associated recognition motif is bipartite (5’-GAAAN_8_TGG-3’), with N6-methyladenine (m6A) occurring at the terminal adenine of the GAAA half-site. Methylation at this motif was readily detected in the WT strain but was absent in the Δ*M. FcoHsdM* mutant (Fig. 1C), indicating that M. FcoHsdM is responsible for this methylation signature in *F. columnare*.

**Figure 1.**
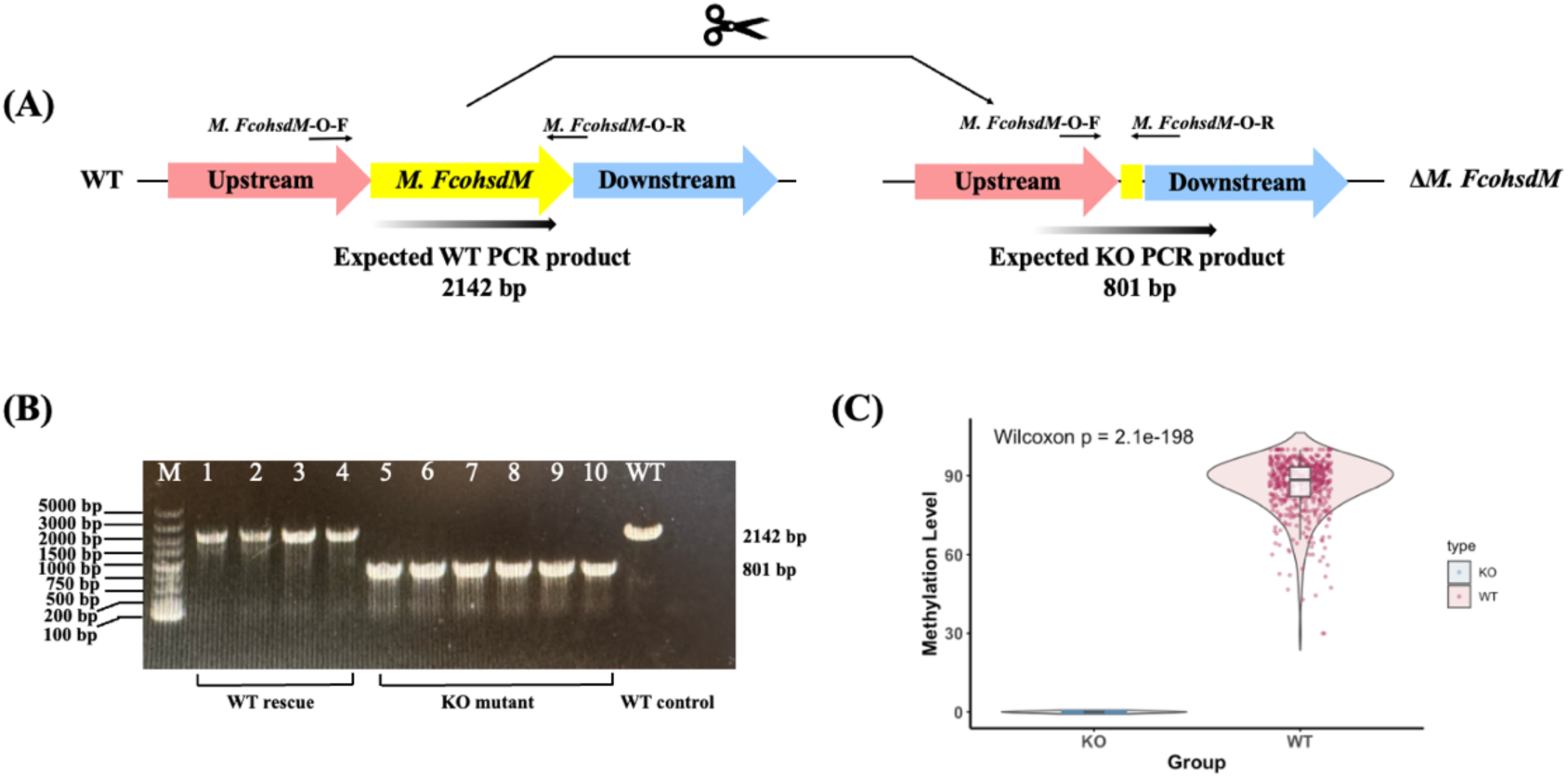
Construction and validation of the *M. FcoHsdM* knockout strain. **(A)** Schematic representation of the *M. FcoHsdM* gene deletion strategy via homologous recombination. The binding sites for the validation PCR primers (*M. FcoHsdM*-O-F and *M. FcoHsdM*-O-R) are indicated by black arrows. The expected sizes of the PCR products are 2142 bp for the wild-type (WT) strain and 801 bp for the Δ*M. FcoHsdM*; **(B)** PCR screening of sucrose-resistant colonies to identify the double-crossover mutants. Lane M: DL 5000 DNA marker; Lanes 1-4: colonies that reverted to the wild-type genotype (2142 bp); Lanes 5-10: positive knockout mutants showing the truncated PCR product (801 bp); Lane WT: wild-type genomic DNA control; **(C)** Comparison of global methylation levels at the *M. FcoHsdM* specific motifs between the Δ*M. FcoHsdM* (KO) and wild-type *F. columnare* (WT) strains. Following *M. FcoHsdM* gene knockout, the corresponding sequences in the Δ*M. FcoHsdM* strain loses methylation modifications.

### M. FcoHsdM is dispensable for planktonic growth but required for virulence in zebrafish

We first examined whether loss of M. FcoHsdM-dependent methylation affects basic physiology. Planktonic growth in modified Shieh (MS) broth at 28 ℃ was indistinguishable between WT, Δ*M. FcoHsdM*, and C-*M. FcoHsdM* over a 54-hour period (Fig. 2A), indicating that M. FcoHsdM is dispensable for growth under standard laboratory conditions.

Despite the absence of a detectable growth defect, deletion of *M. FcoHsdM* caused a profound loss of virulence in the zebrafish infection model. Zebrafish challenged with the wild-type strain developed typical signs of columnaris disease before death, and the wild-type infection group reached 100% mortality by day 14 post-challenge. In contrast, all fish challenged with the Δ*M. FcoHsdM* mutant survived throughout the experiment, resulting in a 100% survival rate that was indistinguishable from the medium-only control group (Fig. 2B). The difference in survival between the wild-type and mutant infection groups was statistically significant by Kaplan-Meier log-rank analysis. *F. columnare* colonies were recovered from deceased fish in the wild-type group, supporting columnaris disease as the cause of mortality.

Together, these data show that M. FcoHsdM is dispensable for planktonic growth but is required for virulence in zebrafish. This separation between normal in vitro growth and complete attenuation in vivo suggested that M. FcoHsdM regulates specific pathogenicity-associated traits rather than general bacterial viability.

**Figure 2.**
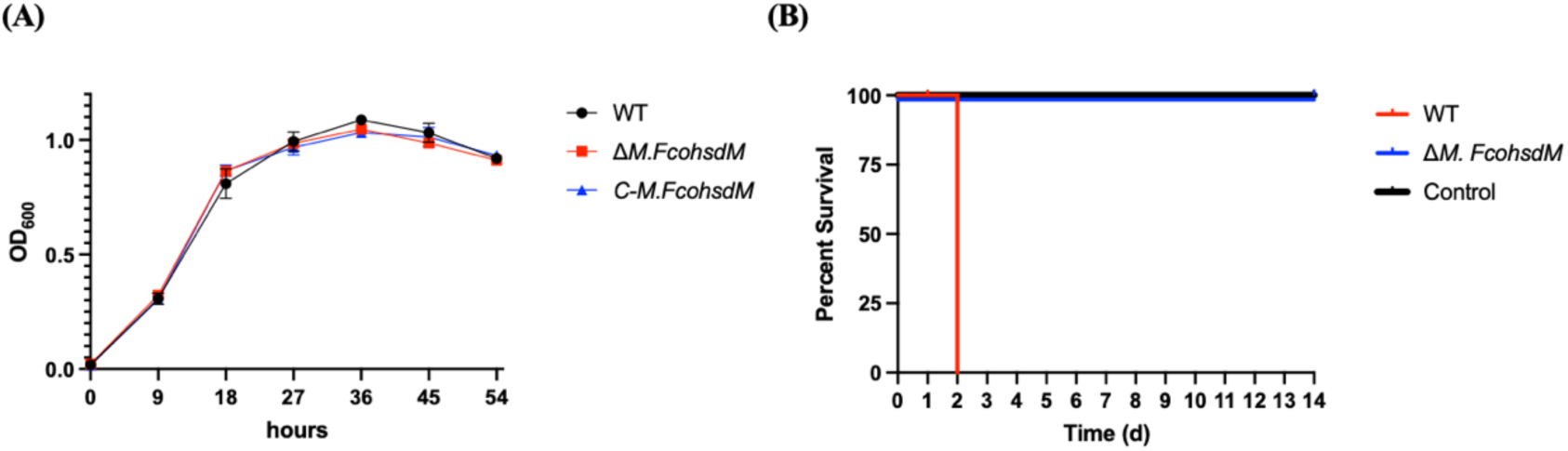
Growth curve analysis and virulence assessment of *F. columnare* wild type and Δ*M. FcoHsdM* mutant. **(A)** Growth curves of the wild-type (WT), the Δ*M. FcoHsdM* mutant, and the complemented strain. Bacteria were cultured in MS broth, and the optical density at 600 nm (OD_600_) was monitored over 54 hours; **(B)** Virulence assessment of *F. columnare* wild type and Δ*M. FcoHsdM* mutant in zebrafish. Three groups of 45 zebrafish (n = 135) were subjected to exposure to the wild type (indicated in red) and the Δ*M. FcoHsdM* mutant (shown in blue). A control group was exposed to an equivalent volume of MS growth medium (shown in black). Survival data were analyzed using Kaplan-Meier log rank survival analysis, and significant differences in percent survival for fish challenged were observed between the wild type and Δ*M. FcoHsdM* (*P*< 0.0001).

### Loss of *M. FcoHsdM* disrupts surface-associated phenotypes and increases OMV production

As *F. columnare* infection depends on surface colonization, spreading motility, and extracellular enzymatic activity, we next examined whether deletion of *M. FcoHsdM* affected phenotypes associated with host interaction and environmental persistence. Biofilm formation was strongly impaired in the Δ*M. FcoHsdM* mutant. Crystal violet staining showed that the mutant produced markedly less biofilm biomass than the wild-type strain after 48 h of incubation in 96-well polystyrene plates. This defect was restored in the complemented strain, indicating that M. FcoHsdM positively regulates biofilm development (Fig. 3A).

The gliding motility was examined, a surface-associated behavior closely linked to colonization and biofilm formation in *F. columnare*. Time-lapse microscopic tracking showed that wild-type cells displayed active surface movement, whereas Δ*M. FcoHsdM* cells were largely stationary or exhibited severely reduced displacement. Complementation restored gliding motility, confirming that the motility defect resulted from loss of *M. FcoHsdM* rather than a secondary mutation (Fig. 3C and D; Movie S1). Deletion of *M. FcoHsdM* also altered colony morphology. When spotted on MS agar and incubated at 28°C, wild-type colonies displayed a yellowish, rough-surfaced morphology, whereas Δ*M. FcoHsdM* colonies appeared smooth. Microscopic examination of individual colonies further showed that wild-type cells formed thin, spreading colonies, while the mutant formed compact, non-spreading colonies (Fig. 3B). These changes are consistent with the observed gliding defect and suggest altered cell-surface properties in the mutant.

In addition to surface colonization traits, we assessed extracellular proteolytic activity. API 20NE biochemical profiling revealed that most tested metabolic and enzymatic traits were unchanged between the wild-type and Δ*M. FcoHsdM* strains. However, the mutant specifically lost gelatinase (GEL) activity, and this defect was restored in the complemented strain (Table S1; Fig. S1A). The loss of proteolytic capacity was further confirmed by the absence of gelatin liquefaction in nutrient gelatin stab assays (Fig. S1B). Antibiotic susceptibility profiles were broadly similar between the wild-type and mutant strains under the tested conditions (Table S2).

Collectively, these phenotypic assays indicate that M. FcoHsdM controls multiple surface and secretion-associated traits, including biofilm formation, gliding motility, colony spreading and gelatinase activity. These defects provide plausible contributors to the loss of virulence observed in zebrafish.

**Figure 3.**
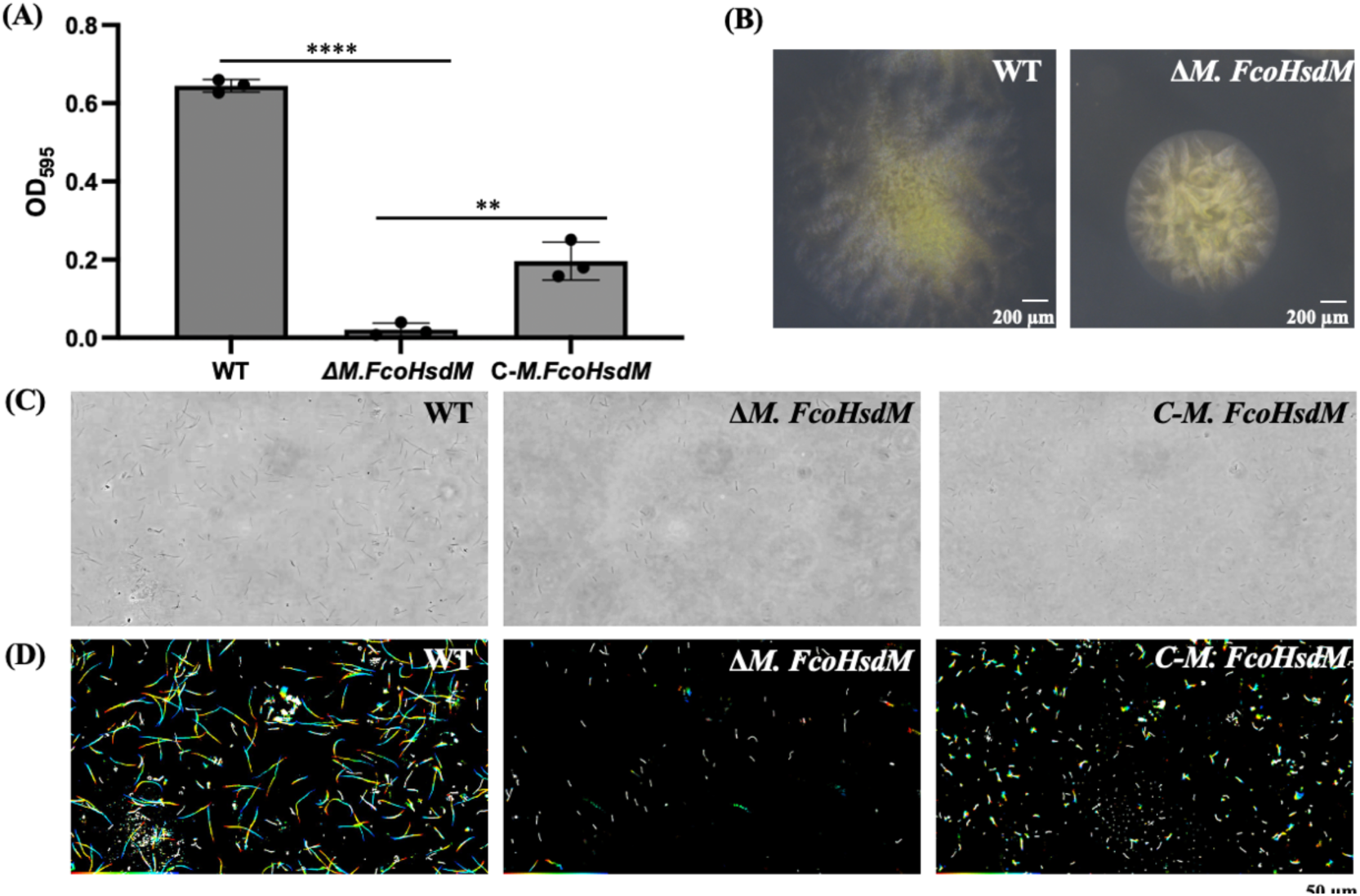
Effects of *M. FcoHsdM* deletion on biofilm formation and gliding motility in *F. columnare*. **(A)** Quantification of biofilm formation. Biofilm biomass was assessed using a crystal violet staining assay in 96-well microplates after 48 hours of incubation. The retained dye was quantified by measuring the absorbance at 595 nm (OD_595_). Data are presented as the mean ± standard deviation (SD) of three independent biological replicates. Significant differences between groups were determined using one-way ANOVA (** *P <* 0.01, **** *P <* 0.0001). **(B)** Photomicrographs of wild-type and mutant strains on MS agar. Colonies were captured using a Nikon SMZ25 microscope after 48 h of incubation on MS agar at 28°C. The image of the WT strain was characterized for spreading colony, whereas the image of the Δ*M. FcoHsdM* mutant formed a non-spreading colony. **(C, D)** Gliding motility of wild-type *F. columnare*, Δ*M. FcoHsdM* mutant, and the complemented strain on glass surfaces. The wild-type *F. columnare*, the Δ*M. FcoHsdM* mutant, and the C-*M. FcoHsdM* strain were grown overnight at 28 °C. For the assay, 10 μl of each bacterial suspension was introduced into glass tunnel slides. Cell motility was captured via an Olympus CKX53 microscope. Single frames from the recordings were temporally color-coded. The color scale started at red (0 seconds) and progressed through orange, yellow, green, and cyan, ending at blue (30 seconds). These colored frames were combined to produce a ‘rainbow trace’ of the gliding cells. **(C)** The initial frame of the motility video for each strain. **(D)** The corresponding 30-second rainbow trajectories. Stationary or minimally moving cells appear white. The 50 μm scale bar applies to all images. The displayed tracks correspond to the dynamic sequences in Movie S1.

### Loss of M. FcoHsdM increases outer membrane vesicle (OMV) production

Transmission electron microscopy (TEM) revealed vesicle-like structures surrounding cells of both strains, with a greater number observed around mutant cells (Fig. 4A). Quantification yielded 8.18 ± 2.87 vesicles per cell for the mutant and 3.84 ± 1.60 vesicles per cell for the WT strain, corresponding to an approximately 2.1-fold increase (Fig. 4B; *P* < 0.001). These observations suggest that loss of M.FcoHsdM alters cell-envelope-associated physiology and increases outer-membrane vesicle production under the conditions tested.

**Figure 4.**
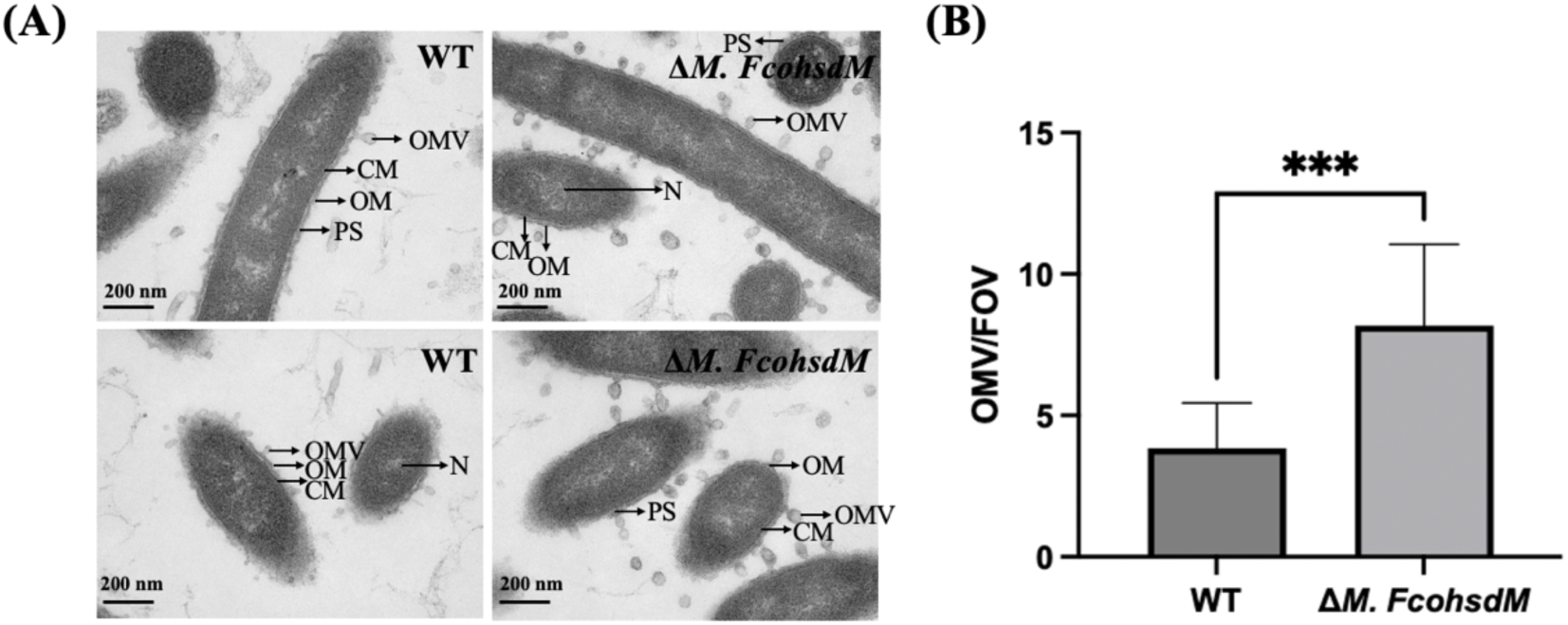
Ultrastructural comparison between the *F. columnare* wild-type and Δ*M. FcoHsdM* strains. **(A)** TEM images of the wild-type (WT) and Δ*M. FcoHsdM* mutant. Cellular structures are highlighted by arrows: N (nucleoid), CM (cell membrane), OM (outer membrane), OMV (outer membrane vesicle), and PS (periplasmic space). The scale bar represents 200 nm; **(B)** OMV quantification based on 12 random fields of view (FOV) per group. Production levels are expressed as the ratio of vesicles to bacterial cells per FOV. Data represent the mean ± SD. Significance was determined via an unpaired Student’s t-test incorporating Welch’s correction.

### Deletion of M. FcoHsdM represses a putative T6SS locus and downregulates secretion-associated genes, including a putative gelatinase

To investigate the molecular basis of the Δ*M. FcoHsdM* mutant phenotypes, we performed RNA-seq on wild-type and mutant strains grown to the early stationary phase in MS medium. Transcriptomic analysis identified 330 differentially expressed genes (DEGs) in the mutant (|Fold change| > 2, FDR < 0.05), consisting of 61 upregulated and 269 downregulated genes (Fig. 5A). Thus, loss of M. FcoHsdM was associated predominantly with transcriptional repression.

Among the most strongly downregulated genes was a putative T6SS locus. Repressed transcripts encoded predicted components of multiple T6SS modules, including baseplate-associated proteins (TssK, TssG, and TssF), a sheath-associated component (TssC), tube/spike-associated proteins (Hcp and VgrG), and a predicted antibacterial effector (Tle1). qRT-PCR analysis of representative T6SS genes confirmed the RNA-seq trends (Fig. 5B), supporting a strong association between M. FcoHsdM and expression of the putative T6SS locus.

In addition to T6SS-associated genes, RNA-seq revealed reduced expression of several genes encoding predicted T9SS-associated proteins. These included genes encoding proteins with type A sorting domains, predicted extracellular metalloproteases, thermolysin-like proteases, glycosyl hydrolases, and gliding motility-associated C-terminal domain-containing proteins. Because T9SS-dependent secretion is required for motility and virulence in *F. columnare*, we validated selected T9SS-associated transcripts by qRT-PCR. The qRT-PCR results were consistent with the RNA-seq data and confirmed that these genes were downregulated in the Δ*M. FcoHsdM* mutant (Fig. 5C). These findings are consistent with the observed defects in gliding motility, gelatin hydrolysis, colony spreading, and virulence, and suggest that altered expression of T9SS-associated genes may contribute to the attenuated phenotype.

Functional enrichment analysis further supported broad changes in surface and secretion-associated physiology. GO terms related to isoprenoid metabolism, cell-envelope functions, and stress responses were enriched among DEGs, while KEGG analysis highlighted bacterial secretion system-related pathways (Fig. 5D and E). These enrichments were consistent with the impaired motility, reduced biofilm formation, and changes in cell-surface structures observed in the Δ*M. FcoHsdM* mutant.

A gene encoding a predicted M4-family metalloprotease, which we designated *fco1*, was also downregulated in the Δ*M. FcoHsdM* mutant. Fco1 shares 40.54% and 31.4% amino acid identity with the well-characterized metalloproteases LasB (*Pseudomonas aeruginosa*) and GelE (*Enterococcus faecium*), respectively [25, 26]. It also exhibits 31.36% identity with Fpp1 from *Flavobacterium psychrophilum*, a protease known to degrade gelatin and host collagens [27]. Further domain analysis using SMART indicated that Fco1 contains an N-terminal FTP (fungalysin/thermolysin propeptide) domain, followed by a central Peptidase_M4 catalytic domain and a C-terminal Peptidase_M4_C domain (Fig. S2A). Within this catalytic region, we identified the conserved HEXXH zinc-binding sequence, which is typical of Zn^2+^-dependent metalloproteases and essential for collagen substrate hydrolysis (Fig. S2B). Methylome data revealed an M. FcoHsdM recognition site located inside the *fco1* coding sequence (CDS), and both fco1 transcripts and protein were reduced in the mutant. Fco1 therefore represents a candidate contributor to the gelatinase-related phenotype observed in the mutant, although direct genetic validation will be required to determine whether it is responsible for gelatin hydrolysis in *F. columnare*.

**Figure 5.**
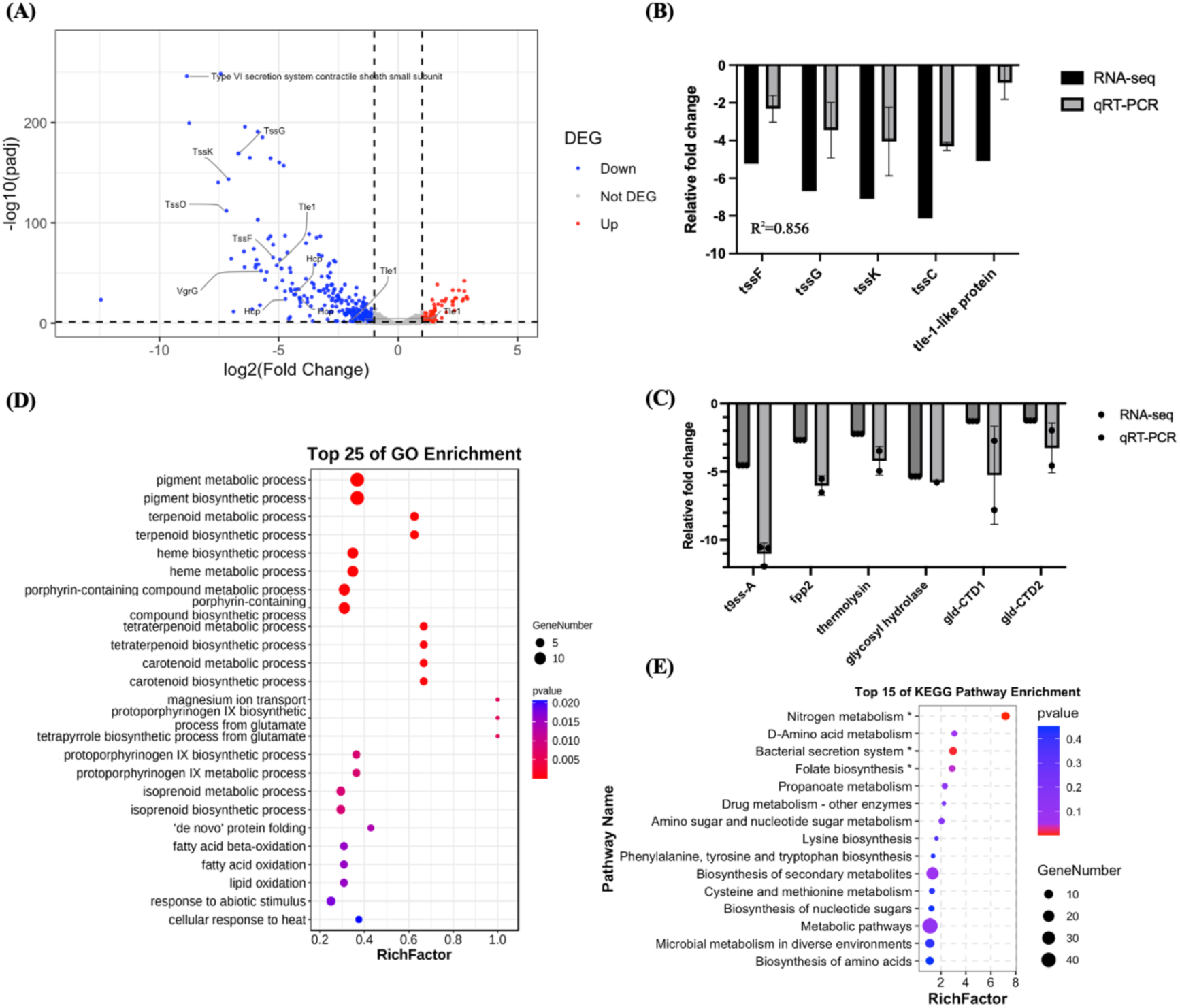
Transcriptomic analysis of the wild-type and Δ*M. FcoHsdM* mutant strains. **(A)** Volcano plot of differentially expressed genes (DEGs) (FDR < 0.05, |log2(fold change)| ≥ 1). Red, blue, and grey dots represent significantly upregulated, significantly downregulated, and non-differentially expressed genes, respectively; **(B)** qRT-PCR validation of selected T6SS-associated genes. The bar chart compares the relative fold changes of selected genes obtained from RNA-seq and qPCR. Error bars represent the standard error of three biological replicates; (C) qRT-PCR validation of selected T9SS-associated genes. Bars indicate RNA-seq fold changes and dots indicate qRT-PCR fold changes for selected genes in the Δ*M. FcoHsdM* mutant relative to the wild-type strain. T9SS-A, T9SS type A sorting domain-containing protein; Fpp2, metalloprotease Fpp2; Gld-CTD1 and Gld-CTD2, gliding motility-associated C-terminal domain-containing proteins 1 and 2. Data are presented as means ± SD from three biological replicates. **(D)** Top 25 enriched GO terms of the DEGs between the WT and Δ*M. FcoHsdM* strains; **(E)** Top 15 enriched KEGG pathways of the DEGs between the WT and Δ*M. FcoHsdM* strains. The asterisk (*) on the y-axis indicates a significant difference.

### M. FcoHsdM-dependent methylation motifs are enriched in coding regions and overlap secretion-associated DEGs

Genome-wide methylation profiling identified 455 M. FcoHsdM-associated motif sites, of which 437 were located within predicted coding sequences. To examine whether these motif-containing genes were associated with transcriptional changes, we compared the methylome and RNA-seq datasets. Sixty-three DEGs overlapped genes containing M. FcoHsdM-associated motifs, and 61 of these motif-containing DEGs carried the motif within coding regions (Fig. 6A). Functional enrichment of motif-containing DEGs suggested associations with metal ion homeostasis, cell-wall polysaccharide metabolism, and defense-related functions (Fig. S3).

Within the putative T6SS locus, *tssP* contained two M. FcoHsdM-associated motifs in its coding sequence. The *tssP* gene is located at the 5′ end of the putative T6SS gene cluster and encodes a predicted membrane-complex component related to Bacteroidota T6SS systems (Fig. 6B). This position makes *tssP* a candidate regulatory entry point for the coordinated downregulation of the T6SS locus observed in the Δ*M. FcoHsdM* mutant. Further sequence analysis using the PRODORIC database (http://prodoric.tu-bs.de/) revealed that a predicted recognition motif for the transcription factor Lrp (Leucine-responsive regulatory protein) is located within the CDS region of *tssP* and directly overlaps with the M. FcoHsdM methylation motif. Given that Lrp is a characterized global regulator previously reported to modulate T6SS expression in other bacterial species [28, 29], this unique structural overlap suggests that Lrp may act as a transcriptional repressor for the *tssP* operon, functioning through an epigenetic switch mechanism. We hypothesize that in the wild-type strain, M. FcoHsdM-mediated DNA methylation sterically hinders Lrp from binding to this site, thereby permitting T6SS expression. Conversely, the loss of methylation in the Δ*M. FcoHsdM* mutant allows Lrp to bind to the unmethylated motif, leading to the repression of *tssP* and the downstream T6SS cluster. Although this proposed interplay between DNA methylation and Lrp binding provides a compelling mechanistic model, it requires further experimental validation.

**Figure 6.**
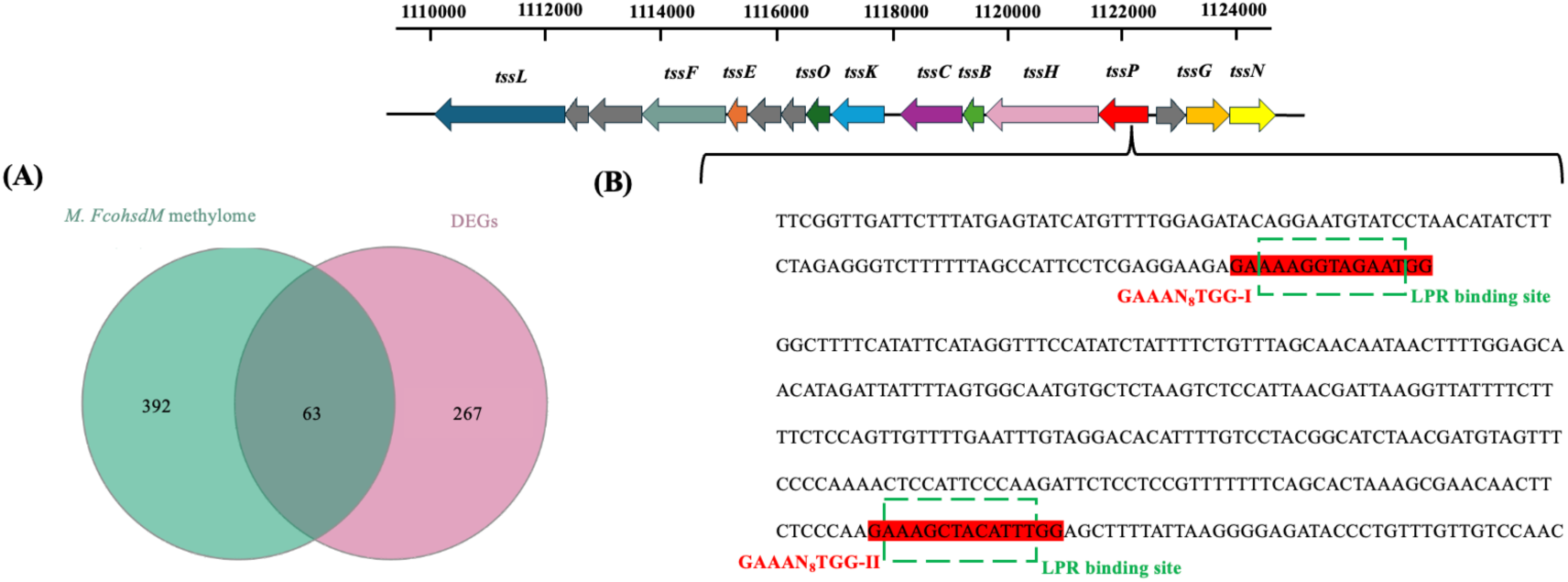
Identification of *tssP* as a direct target of M. FcoHsdM methylation within the T6SS gene cluster. **(A)** A Venn plot shows the overlapped genes between DEGs and genes within the M. FcoHsdM motif identified in the Δ*M. FcoHsdM* strain. A total of 63 overlapping genes were identified; **(B)** Genomic organization of the T6SS gene cluster and sequence analysis of the *tssP* gene. The upper panel shows the arrangement of genes within the T6SS cluster, with *tssP* (highlighted in red) positioned as the first gene of the transcriptional unit. The lower panel displays a partial nucleotide sequence of the *tssP* CDS region. The two identified M. FcoHsdM methylation motifs (GAAAN_8_TGG-I and GAAAN_8_TGG-II) are highlighted in red boxes. The predicted recognition sites for the transcription factor Lrp are indicated by green dashed boxes, demonstrating direct overlap with the methylation motifs.

### Extracellular proteomic analysis supports altered secretion-associated output in the Δ*M. FcoHsdM* mutant

To determine whether transcriptional changes were reflected at the protein level, we performed LC-MS/MS analysis of extracellular protein fractions from WT and Δ*M. FcoHsdM* cultures. A total of 1,636 proteins were identified, among which 516 showed significant differences between the two strains, including 444 increased and 72 decreased proteins in the mutant (absolute fold change ≥ 1.5, Benjamini-Hochberg-adjusted *P* < 0.05) (Figure 7A and 7B). We focused on proteins associated with secretion, envelope functions, motility, and T6SS-related pathways.

Consistent with the RNA-seq results, several T6SS-associated proteins showed reduced abundance in the Δ*M. FcoHsdM* mutant, including Hcp, TssC, the type VI secretion system contractile sheath small subunit, and MepH (Fig. 7C). In contrast, selected T9SS-, gliding motility-, and surface-associated proteins showed more heterogeneous changes. Several proteins, including GldJ, GldL, GldG, and an Ig-like domain-containing protein, were increased in the mutant, whereas multiple predicted extracellular enzymes or T9SS-associated proteins, including Fco1, a thermolysin-like protein, Fpp2, glycosyl hydrolase, and proteins containing T9SS C-terminal sorting or target domains, were decreased (Fig. 7C). These data suggest that deletion of M. FcoHsdM does not simply increase or decrease extracellular protein abundance globally, but instead reshapes secretion and surface associated protein output in a pathway and protein specific manner.

**Figure 7.**
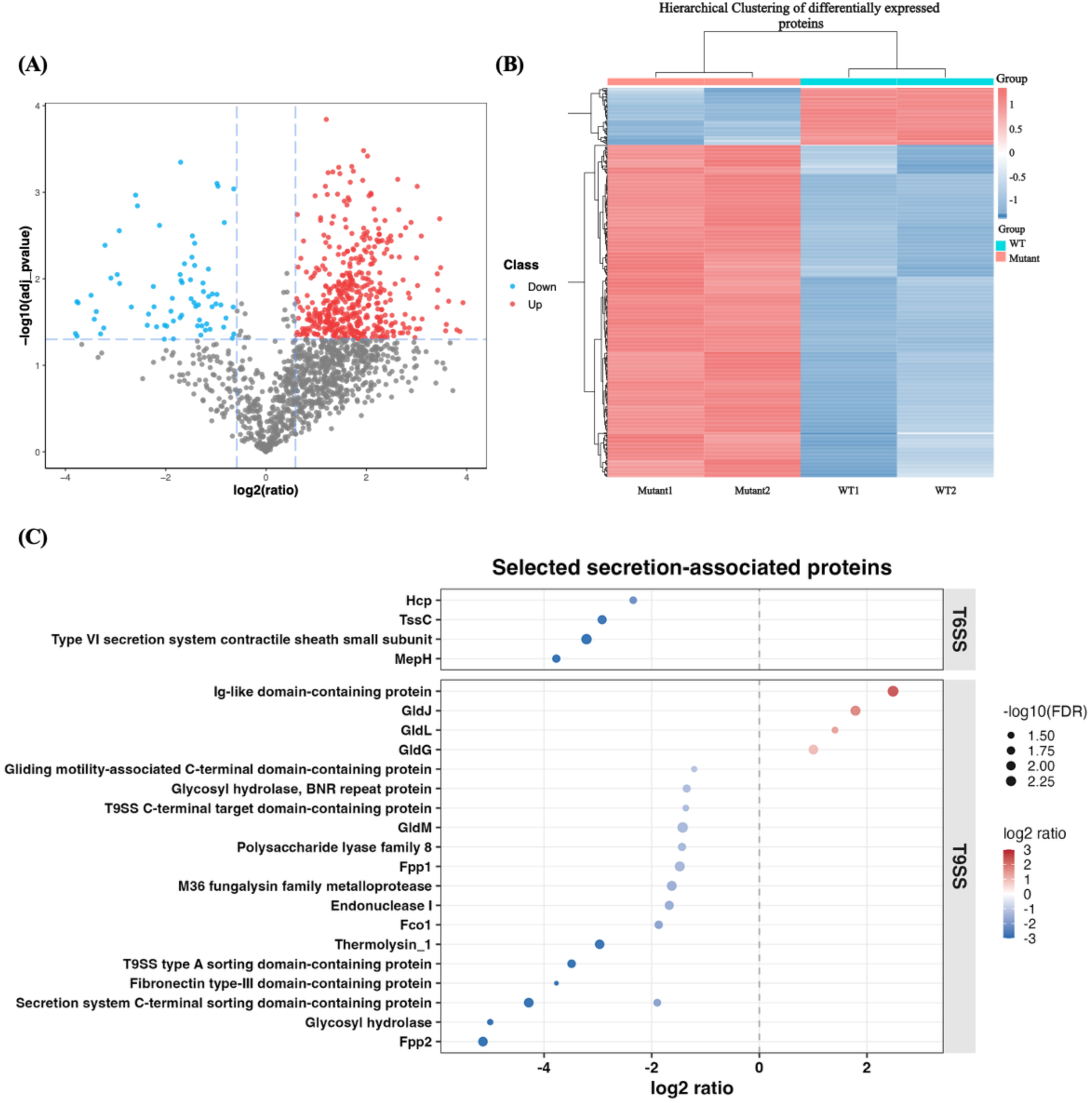
Proteomic analysis for differentially expressed genes. (**A)** Volcano map of differential genes in proteomics (FDR < 0.05, |fold change| ≥ 1.5). The red dot represents significantly upregulated differences; the blue dot represents significantly downregulated differences; the grey dot represents no difference; **(B)** Heat map of the two-way hierarchical clustering; **(C)** Bubble plot of selected secretion associated proteins. Dot color represents the log_2_-transformed abundance ratio of Δ*M. FcoHsdM* relative to WT, and dot size represents -log_10_-transformed FDR. Selected proteins were grouped into T6SS- and T9SS-associated categories based on annotation. Blue indicates lower abundance and red indicates higher abundance in the Δ*M. FcoHsdM* mutant relative to WT.

### Integrated analysis revealed the global roles of *M. FcoHsdM* in *F. columnare*

To determine whether the transcriptional changes caused by loss of M. FcoHsdM are propagated to the protein level, we compared the transcriptome and proteome of the Δ*M. FcoHsdM* and WT strains. The transcriptomic and proteomic profiles identified 2902 mRNAs and 1524 proteins, respectively, of which 1521 identified proteins were assigned to the identified mRNAs. Of all the DEGs identified in the Δ*M. FcoHsdM* strain, 168 were included in the 1521 genes identified in both the proteomes and transcriptomes (Fig. 8A and 8B). As shown in Figure 8C, 783 genes were significantly affected at the mRNA and/or protein levels in the Δ*M. FcoHsdM* strain. Among the 783 genes, 453 were found to be significantly affected only at the protein level (NoDE_Genes and DE_Prots), 105 were significantly affected only at the mRNA level (DE_Genes and NoDE_Prots), and 63 were significantly affected at both the mRNA and protein levels (DE_Genes and DE_Prots) (Fig. 8C). Among the 63 co-differentially expressed genes, 53 showed consistent expression trends (both up-or down-regulated), while 10 showed opposite trends (Fig. 8D). Additionally, 162 genes were differentially expressed only at the mRNA level but were not detected in the proteome. GO enrichment analysis revealed that these co-differentially expressed genes were primarily involved in biological processes such as proteolysis, cellular glucan metabolic process, and vesicle-mediated transport (Fig. 8E). To elucidate the functional relationships among these significantly altered candidates, 63 DEPs that overlap with the identified DEGs were shown to be involved in a Protein-protein interaction (PPI) network based on the STRING database (score > 700). Notably, a prominent module consisted of core components of the T6SS, including TssB, TssC, and TssP, along with MepH. All nodes within this cluster exhibited consistent downregulation at both the mRNA and protein levels. This indicates that *M. FcoHsdM* is critical for coordinating the T6SS apparatus, which is closely related to interbacterial competition. Furthermore, another tightly linked module comprised key molecular chaperones and heat shock proteins, specifically ClpB, GrpE, and GroES. The concerted alteration of these stress-response proteins suggests that the deletion of *M. FcoHsdM* disrupts intracellular protein homeostasis or induces cellular stress, thereby triggering a robust chaperone-mediated compensatory response (Fig. 8F).

**Figure 8.**
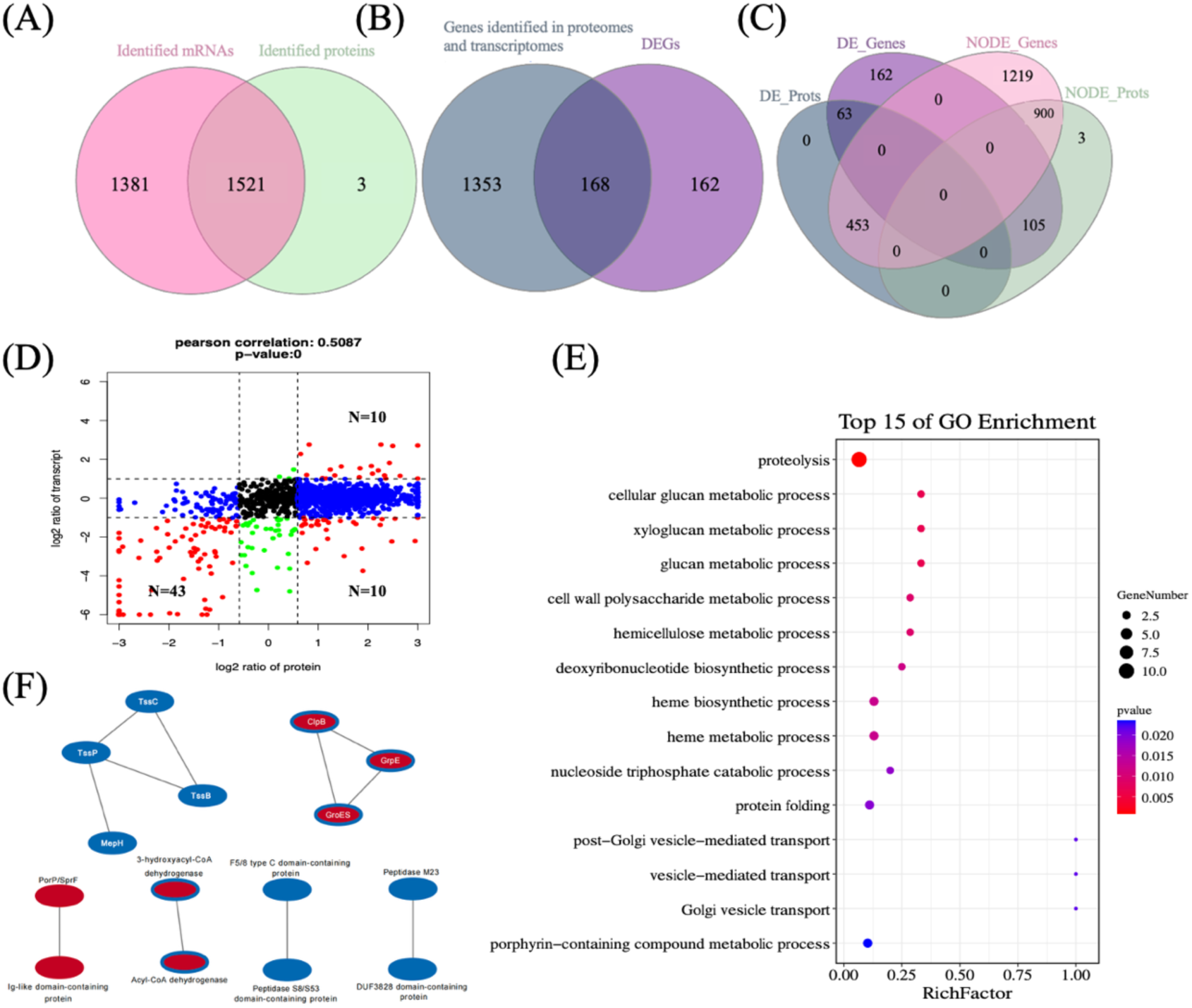
Integrated proteomic and transcriptomic analyses. (**A)** Venn diagram showing that the 1521 proteins identified in the proteomes can be assigned to the 2902 mRNAs identified in the transcriptomes. The pink circle indicates all mRNAs identified in both Δ*M. FcoHsdM* and WT; the green circle indicates all proteins identified in both Δ*M. FcoHsdM* and WT; (**B)** Venn diagram showing that 168 DEGs were included in the 1521 genes identified in both the proteomes and transcriptomes. The purple circle indicates DEGs in Δ*M. FcoHsdM*, the blue circle indicates 1521 genes identified in both the proteomes and transcriptomes; (**C)** Venn diagram showing the effects of *M. FcoHsdM* deletion on wild-type strain at the mRNA and/or protein levels. The blue ellipse indicates proteins significantly affected by *M. FcoHsdM* deletion in the proteomic profiles; the purple ellipse indicates genes significantly affected by *M. FcoHsdM* deletion in the transcriptomic profiles; the pink ellipse indicates genes not affected by *M. FcoHsdM* deletion in the transcriptomic profiles; the green ellipse indicates proteins not affected by *M. FcoHsdM* deletion in the proteomic profiles; **(D)** Nine-quadrant scatter plot showing the correlation of expression changes between the transcriptome and proteome. The x-axis and y-axis represent the log2 fold changes of proteins and transcripts, respectively; **(E)** Bubble chart showing the top 15 enriched GO terms of the differentially expressed candidates. The size of the circles represents the number of genes, and the color indicates the p-value; **(F)** PPI network of the significantly altered candidates. Distinct functional modules, such as the T6SS apparatus and molecular chaperones, are highlighted.

### M. FcoHsdM promotes T6SS-associated interbacterial competition

To determine whether the transcriptional repression of the T6SS cluster in the Δ*M. FcoHsdM* mutant was associated with impaired antibacterial activity, we performed a solid-surface interbacterial competition assay using *F. columnare* as the predator and *Aeromonas hydrophila* ATCC 7966 as the prey. In the prey-alone control, *A. hydrophila* maintained high viability after incubation. Co-incubation with wild-type *F. columnare* markedly reduced the recovery of viable *A. hydrophila*, whereas co-incubation with the Δ*M. FcoHsdM* mutant resulted in significantly higher prey survival than the wild-type competition group (Fig. 9). These results indicate that deletion of M. FcoHsdM compromises *F. columnare*-mediated interbacterial antagonism. Together with the transcriptomic and proteomic downregulation of T6SS components, this finding provides functional evidence that M. FcoHsdM contributes to T6SS-associated interbacterial competition.

**Figure 9.**
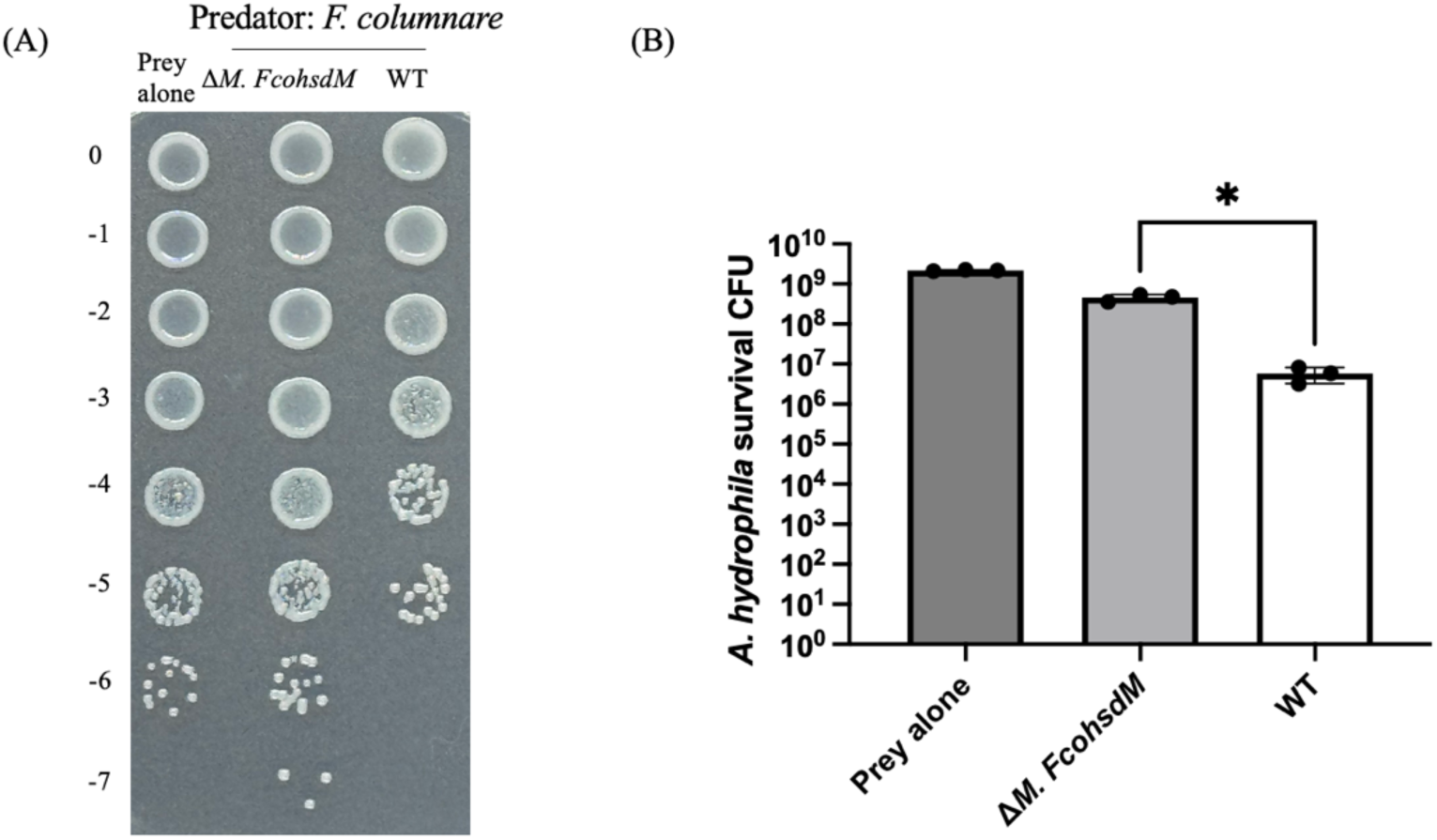
Loss of M. FcoHsdM weakens interbacterial killing by *F. columnare*. **(A)** Representative serial dilution spot assay showing survival of *A. hydrophila* ATCC 7966 after competition with *F. columnare* WT or Δ*M. FcoHsdM*. The prey-alone control contained *A. hydrophila* without *F. columnare*. Numbers indicate 10-fold serial dilutions. **(B)** Competition assay using *F. columnare* WT and Δ*M. FcoHsdM* as predators and *A. hydrophila* ATCC 7966 as prey. Bar graphs represent the means ± SD from three biological replicates. Statistical analyses were performed using the unpaired Student’s t test; \**P* < 0.01.

## DISCUSSION

DNA methylation by restriction–modification systems has traditionally been viewed as a defense mechanism that protects bacterial genomes from foreign DNA [30]. Increasing evidence, however, indicates that bacterial methyltransferases can also act as epigenetic regulators of transcription, phase variation, stress adaptation, and virulence [20, 31]. In this study, we characterized the type I methyltransferase M.FcoHsdM in *F. columnare* and showed that it is required for a major methylation signature in this pathogen. Deletion of *M.FcoHsdM* did not impair planktonic growth under standard laboratory conditions, but caused marked defects in pathogenicity-associated phenotypes, including reduced biofilm formation, impaired gliding motility, loss of gelatinase activity, altered outer membrane vesicle production, attenuated virulence in zebrafish, and reduced interbacterial antagonism. Together with transcriptomic, methylomic, and extracellular proteomic analyses, these results support a model in which M.FcoHsdM-dependent methylation contributes to the regulation of surface and secretion-associated functions in *F. columnare*.

A prominent phenotype of the Δ*M.FcoHsdM* mutant was the disruption of cell-surface-associated behaviors. The mutant formed less biofilm, exhibited defective gliding motility, and produced compact non-spreading colonies. These phenotypes are highly relevant to *F. columnare* pathogenesis, because surface attachment, spreading motility, and colonization of external fish tissues are central features of columnaris disease [2]. The transcriptomic data provide candidate molecular explanations for these defects. Genes associated with gliding motility, cell-envelope functions, extracellular enzymes, and predicted T9SS-associated proteins or substrates were altered in the mutant. Because the T9SS is required for secretion of motility adhesins, enzymes, and other surface-exposed proteins in members of the Bacteroidota [32–34], altered expression or extracellular abundance of T9SS associated proteins could contribute to the observed defects in motility, biofilm formation, colony morphology, and virulence [4]. However, the proteomic changes among T9SS/surface-associated proteins were not uniformly directional, indicating that loss of M.FcoHsdM reshapes extracellular protein output rather than simply suppressing the entire T9SS pathway.

The loss of gelatinase activity in the Δ*M.FcoHsdM* mutant provides another potential explanation for its attenuation. Gelatinase and extracellular metalloproteases can promote bacterial invasion and tissue damage by degrading host structural proteins and extracellular matrix components [5, 35, 36]. In this study, the mutant specifically lost gelatinase activity in biochemical assays, and a candidate M4-family metalloprotease, Fco1, was reduced at both the transcript and protein levels. Fco1 is annotated as a putative extracellular metalloprotease based on its conserved catalytic domain architecture. Moreover, the presence of an M.FcoHsdM-associated methylation motif within the fco1 coding sequence suggests that Fco1 may be connected to the M.FcoHsdM-dependent regulatory network. Nevertheless, direct genetic evidence is still required to determine whether Fco1 is responsible for gelatin hydrolysis and whether methylation at the fco1 locus directly controls its expression. Construction of an fco1 deletion mutant and complementation strain will be important for testing this possibility.

One of the strongest molecular signatures of the Δ*M.FcoHsdM* mutant was repression of a putative T6SS locus. RNA-seq revealed reduced expression of genes encoding predicted T6SS baseplate, sheath, tube/spike, membrane-associated, and effector components, and extracellular proteomics confirmed reduced abundance of several T6SS-associated proteins, including Hcp, TssC, TssP-related components, and MepH. Integrated transcriptomic and proteomic analyses further identified a concordantly downregulated T6SS module. Importantly, the interbacterial competition assay provided functional support for these omics results: the Δ*M.FcoHsdM* mutant showed reduced antagonistic activity against *A. hydrophila* compared with the wild-type strain. These findings indicate that M.FcoHsdM contributes to T6SS-associated antibacterial competition in *F. columnare*. Because bacterial competition can influence pathogen fitness in polymicrobial aquatic environments and on host mucosal surfaces, reduced T6SS activity may affect ecological competitiveness and infection dynamics [37–39].

The methylome analysis suggests a possible mechanism linking M.FcoHsdM-dependent methylation to T6SS regulation. Most M.FcoHsdM-associated motifs were located within coding regions, and a subset of motif-containing genes overlapped with differentially expressed genes. Within the putative T6SS locus, *tssP* contained M.FcoHsdM-associated motifs in its coding sequence. This is particularly interesting because *tssP* lies near the 5′ region of the predicted T6SS gene cluster and encodes a predicted Bacteroidota-specific membrane complex component. Sequence analysis further identified a predicted Lrp-binding motif overlapping one of the M.FcoHsdM-dependent methylation motifs. This arrangement raises the possibility that methylation within the *tssP* gene body modulates transcription factor occupancy and thereby influences expression of the T6SS locus. Such a model is conceptually consistent with established bacterial epigenetic switches in which DNA methylation affects regulator binding, although the genomic context here is distinct because the candidate methylation site lies within a coding sequence rather than a canonical promoter-proximal region.

Based on these observations, we propose a working model in which M.FcoHsdM-dependent methylation helps maintain expression of the putative T6SS locus, potentially by interfering with binding of a regulatory protein such as Lrp at the *tssP* coding region. In the Δ*M.FcoHsdM* mutant, loss of methylation may increase access of this regulator to the unmethylated site, leading to repression of *tssP* and downstream T6SS-associated genes. This observation aligns with the emerging paradigm of intragenic DNA methylation in eukaryotes, where gene body methylation is thought to maintain transcriptional continuity by inhibiting H2A.Z deposition[40], or enhance transcriptional efficiency by suppressing spurious initiation from alternative promoters [41, 42]. We hypothesize that in *F. columnare*, M. FcoHsdM-mediated gene body methylation may exert similar effects through two distinct molecular mechanisms. First, methylation could act via direct steric hindrance, as exemplified by the Lrp interaction. The addition of methyl groups may physically block Lrp from binding to its target DNA, thereby altering local DNA topology and RNA polymerase processivity. Alternatively, methylation might function through an indirect, synergistic mechanism to suppress non-specific transcriptional initiation. This process could involve specific cofactors that recognize and bind to conserved DNA motifs. The methylation status of these genomic elements could modulate the recruitment or activity of these cofactors, which subsequently interact with transcription factors to co-regulate gene expression. However, this model remains hypothetical and requires direct experimental validation.

In summary, our findings identify M.FcoHsdM as an important methylation-dependent regulator of pathogenicity-associated traits in *F. columnare*. Loss of M.FcoHsdM-dependent methylation leads to broad changes in surface behavior, extracellular protease activity, OMV production, secretion-associated gene expression, and T6SS-mediated interbacterial competition. The discovery of M.FcoHsdM-associated methylation motifs within coding regions, including the *tssP* locus, suggests that gene-body methylation may contribute to bacterial transcriptional regulation by modulating local regulator-DNA interactions. These findings expand our understanding of type I restriction–modification methyltransferases as regulatory factors in aquatic bacterial pathogens and provide a framework for future studies on methylation-controlled secretion systems, virulence, and microbial competition in *F. columnare*.

## MATERIALS AND METHODS

### Bacterial strains, plasmids, and growth conditions

#### Construction of *M. FcoHsdM* deletion mutant

The bacterial strains, plasmids, and primers used in this study are listed in Tables S3 and S4. A 1880 bp upstream fragment of *M. FcoHsdM* was PCR-amplified from *F. columnare* genomic DNA with primers L-F (KpnI site) and L-R, and a 1301 bp downstream fragment was amplified with R-F and R-R (BamHI site) (Fig. 1A). The two fragments were assembled into KpnI/BamHI-linearized pMS75 using the ClonExpress Ultra One Step Cloning Kit V2 (Vazyme), yielding plasmid pRX03. Plasmid pRX03 was introduced into the *F. columnare* wild type by conjugation, and integrants were selected on tetracycline. Tetracycline-resistant colonies were streaked on MS agar with tetracycline, then cultured in MS broth without tetracycline to promote plasmid loss via recombination. Cultures were plated on MS agar containing 10% sucrose to counterselect the *sacB* marker, allowing growth only of recombinants that had lost the plasmid. Colonies growing on sucrose were screened by PCR to confirm the *M. FcoHsdM* deletion, and the mutant was designated Δ*M. FcoHsdM*.

#### Complementation of the *M. FcoHsdM* gene in Δ*M. FcoHsdM*

A 2361 bp *M. FcoHsdM* fragment was PCR-amplified with primers P23-M-F (BamHI site) and P23-M-R (SphI site) (Table S4), ligated into the shuttle vector pCP23 to generate pCP23-*M. FcoHsdM*, and introduced into the Δ*M. FcoHsdM* by conjugation.

#### Biochemistry and antibiotic resistance test

For biochemical characterization, the API 20NE test system (bioMérieux, France) was used according to the manufacturer’s instructions to identify any biochemical differences between the mutant and wild-type strains. Antimicrobial susceptibility was evaluated using the standard Kirby-Bauer disk diffusion method in accordance with the Clinical and Laboratory Standards Institute (CLSI) guidelines [43]. Briefly, bacterial suspensions were adjusted to an OD600 of 0.5 and uniformly swabbed onto MS agar plates. Commercially available antibiotic susceptibility disks from Hangzhou Microbial Reagent Company (Zhejiang, China) were then placed on the surface of the agar. After incubation at 28 °C for 24 h, the diameters of the zones of inhibition were measured in millimeters to compare the antibiotic resistance profiles of the wild-type and Δ*M. FcoHsdM*.

#### Growth curves

The frozen stocks of *F. columnare* strains were revived by plating on modified Shieh (MS) agar and incubating at 28 °C for 36 hours [44]. Colonies obtained were used to inoculate 10 mL of MS medium, followed by overnight incubation at 28 °C with shaking at 125 rpm. For plasmid-harboring strains, the medium was supplemented with 2.5 μg/mL tetracycline. Overnight cultures of the wild-type, Δ*M. FcoHsdM* mutant and complemented strain were adjusted to an OD_600_ of 0.5 and used to inoculate 10 mL of fresh MS broth. All cultures were incubated at 28 °C with shaking at 125 rpm, and OD_600_ readings were taken every 9 hours over a 54-hour period using a SpectraMax iD3 Multi-Mode Microplate Reader (Molecular Devices, San Jose, CA, USA).

#### Biofilm formation

Biofilm formation in the wild-type and mutant strains was evaluated following an established protocol [45]. Briefly, mid-logarithmic phase cultures (OD₆₀₀ = 0.5) grown in MS medium were diluted 1:100 in fresh MS medium, and 150 μL aliquots were transferred into wells of a 96-well flat-bottom polystyrene microplate (Corning 167008, Corning, NY). The plate was wrapped in aluminum foil and incubated at 28 °C for 48 h. Each strain was tested in triplicate, with sterile uninoculated medium included as a negative control. After incubation, the medium was removed, and the wells were gently washed three times with 200 μL of sterile distilled water. The attached biofilm was stained with 150 μL of 1% (w/v) crystal violet for 30 min at room temperature. Unbound dye was removed by four washes with 200 μL sterile distilled water. The bound dye was subsequently solubilized with 100 μL ethanol, and the absorbance at 595 nm (OD₅₉₅) was recorded using a SpectraMax iD3 Multi-Mode Microplate Reader (Molecular Devices, USA). The final absorbance value for each strain was obtained by subtracting the value of the negative control.

#### Colony morphology assay

To examine individual colony morphology, mid-logarithmic-phase cultures were subjected to 10-fold serial dilutions in sterile MS broth, and 100 μL of appropriate dilutions was plated onto MS agar. After incubation at 28°C for 48 h, individual colonies were observed and photographed using a Nikon SMZ25 stereomicroscope (Nikon, Tokyo, Japan).

#### Cell motility analysis

To evaluate the gliding behavior of individual cells, the cultures were incubated overnight in 1/10 MS broth at 28 °C with constant shaking before observation. Tunnel slides were prepared according to a prior method [46] by combining glass microscope slides, coverslips, and double-sided tape. After preparation, a 10 μl of the culture was introduced into the tunnels. The slides were incubated for 5 minutes before imaging. The cell motility was recorded at 25 °C using an inverted microscope (Olympus CKX53, Tokyo, Japan) equipped with a CMOS camera (Axiocam 208 color, Zeiss, Oberkochen, Germany). Finally, the movement paths were visualized as rainbow traces using the Color FootPrint macro function within Fiji (version 2.14.0) [47].

#### Transmission electron microscopy (TEM) analysis

For TEM observation, overnight bacterial cultures were pelleted by centrifugation and immediately fixed with a TEM-grade fixative at 4°C. After washing three times with 0.1 M phosphate buffer (PB, pH 7.4), the cell pellets were resuspended in 1% agarose and allowed to solidify into blocks. These agarose blocks were then post-fixed using 1% osmium tetroxide (OsO₄) in 0.1 M PB for 2 hours at room temperature in the dark, and rinsed three times with the same buffer. Dehydration was carried out at room temperature using a graded ethanol series (30%, 50%, 70%, 80%, 95%, and 100%), followed by two washes in pure acetone. For resin infiltration, the samples were sequentially incubated in acetone/EMBed 812 resin mixtures at 37°C. The samples were then embedded in pure resin, kept at 37°C overnight, and finally polymerized at 60°C for at least 48 hours. Ultrathin sections (60-80 nm) were cut with an ultramicrotome, mounted onto formvar-coated 150-mesh copper grids, and double-stained with 2% uranyl acetate and 2.6% lead citrate. The prepared grids were then examined and imaged by TEM. For the quantification of OMV production, TEM micrographs from 12 random FOV were analyzed for each strain. The relative OMV production was determined by calculating the ratio of the total number of OMVs to the total number of bacterial cells within each FOV. Data were presented as mean ± standard deviation (SD). Statistical significance was analyzed using an unpaired Student’s t-test with Welch’s correction.

#### Gelatin hydrolysis test

Gelatinase activity was determined using the standard nutrient gelatin stab method. Briefly, fresh bacterial stock (18 h culture) was stabbed into tubes containing nutrient gelatin medium (10% w/v) using a sterile inoculating loop. Following incubation at 28°C for 48 h, the tubes were chilled at 4°C for 30 min. The assay was interpreted immediately upon removal from the refrigerator. An uninoculated tube was included as a negative control.

#### Interbacterial competition assay

Interbacterial competition assays in solid medium were performed as previously described [48]. Briefly, *F. columnare* WT and Δ*M. FcoHsdM* were used as predator strains and *Aeromonas hydrophila* ATCC 7966 as the prey strain. Predator and prey overnight cultures were pelleted, washed three times in fresh MS broth, and resuspended at an OD_600_ of 1.0. Predator and prey cells were mixed at a ratio of 5:1, and 10μl of drops was spotted on an MS agar plate. A prey-alone group containing only *A. hydrophila* was included as the control. After incubation at 28°C for 4 hours, bacterial spots were harvested and resuspended in 0.7mL of sterile MS. The suspensions were serially diluted and plated on ampicillin MS agar plates to determine the number of surviving *A. hydrophila*. CFUs were counted after incubation at 28°C for 24 hours. Three independent biological replicates were performed.

#### Virulence test in zebrafish

Wild-type *F. columnare* and the Δ*M. FcoHsdM* mutant strains were grown overnight in MS broth at 28°C for 24 hours. On the following day, 100 μL of each overnight culture was inoculated into 10 mL of fresh MS medium until the OD₆₀₀ reached approximately 0.4. To determine viable cell counts, serial dilutions were prepared in triplicate and plated on MS agar for enumeration. No symptoms of disease were observed before the challenge, and no *F. columnare* was detected in either the acclimation tanks or the uninfected control groups throughout the experiment.

Challenge assays were performed using triplicate 2-liter containers with restricted water flow at 28°C, each holding 15 fish. Fish were placed into the challenge aquaria one week in advance to allow for acclimatization. Zebrafish were exposed to 600 mL of water containing the bacterial strains at 28°C for 2 hours. After the immersion period, 1.2 L of fresh water was added to reduce the bacterial load and maintain a lower-level continuous exposure. Survival was monitored daily for 14 days. Strains examined were wild type, Δ*M. FcoHsdM*, and a control group. The final challenge concentrations were 1.78 × 10^7^ CFU/mL for wild type, and 1.65 × 10^7^ CFU/mL for Δ*M. FcoHsdM* as quantified by serial dilution. “Control” indicates fish exposed to an equivalent amount of MS growth medium instead of *F. columnare*. Mortalities were monitored, removed, and recorded daily. Data from the triplicate containers for each strain were combined, and survivor fractions were calculated. To verify *F. columnare* infection, the deceased fish were randomly selected and subjected to bacterial examination. Swabs from both external and internal organs were streaked on MS agar, and colonies showing yellow pigmentation, rhizoid, and adherent were identified as *F. columnare*. The present experiment was conducted in compliance with the animal research guidelines of the Hong Kong Special Administrative Region (HKSAR) under animal license [Ref No.: (24-50) in DH/HT&A/8/2/5 Pt.14] and with approval from the City University Animal Ethics Committee (Approval No.: A-0402).

#### RNA-seq

*F. columnare* wild type, Δ*M. FcoHsdM* mutant were grown in MS at 28 °C for 24 h. Each culture was standardized to an OD_600_ of 0.5, and 100 μL was inoculated into 10 mL of fresh MS medium to the early stationary phase. Total RNA was extracted from three biological replicates per strain using the RNeasy Mini Kit (QIAGEN), followed by Dnase I treatment to remove genomic DNA contamination. RNA integrity and concentration were verified using a NanoDrop spectrophotometer and an Agilent 2100 Bioanalyzer. RNA-seq libraries were prepared by Novogene (Beijing, China), involving ribosomal RNA depletion and Illumina-compatible library construction. The pooled libraries were sequenced on an Illumina NovaSeq X Plus platform to generate 150 bp paired-end reads. Raw sequencing data were subjected to quality control using FastQC (v0.11.8) [49], and adapters as well as low-quality bases were trimmed with Trimmomatic (v0.32) [50]. The resulting high-quality reads were aligned to the *F. columnare* reference genome (GCF_049561265.1) using Bowtie2 [51]. Read counts for each gene were generated with FeatureCounts (v2.0.3) [52]. Differential expression analysis was performed using the DESeq2 package (v1.34.0) [53], with genes exhibiting an absolute |log₂(fold change)| > 1 and a false discovery rate (FDR) < 0.05 considered statistically significant [44]. Gene ontology (GO) and Kyoto Encyclopedia of Genes and Genomes (KEGG) enrichment analyses were performed on differentially expressed genes (DEGs). Raw sequencing reads were deposited in the NCBI Sequence Read Archive (SRA) as part of BioProject PRJNA1527449.

#### Quantitative real-time PCR validation

RNA-seq results were validated by quantitative real-time PCR (qPCR) using representative DEGs. Total RNA was isolated with the RNeasy Mini Kit (QIAGEN) and quantified on a NanoDrop 2000 spectrophotometer (Thermo Fisher). Reverse transcription was performed using HiScript IV All-in-One Ultra RT SuperMix (Vazyme) according to the manufacturer’s instructions. qPCR was conducted on a QuantStudio 7 Pro system (Applied Biosystems, Thermo Fisher) using Taq Pro Universal SYBR qPCR Master Mix (Vazyme) and gene-specific primers (Table S4). Thermal cycling conditions comprised initial denaturation at 95 °C for 30 s, followed by 40 cycles of 95 °C for 10 s and 60 °C for 30 s, with a final melt curve stage (95 °C for 15 s, 60 °C for 60 s, and 95 °C for 15 s). Target gene expression was normalized to *gap1* and calculated via the 2^−ΔΔCT^ method [45].

#### DNA methylation sequencing and analysis

Genomic DNA was extracted from *F. columnare* WT and Δ*M. FcoHsdM* strains using the TaKaRa MiniBEST Universal Genomic DNA Extraction Kit (TaKaRa Bio Inc., Japan). For whole-genome sequencing (WGS) and methylation profiling, 1 µg of high-quality native DNA was used for library construction with the Native Barcoding Kit 24 V14 (SQK-NBD114.24, Oxford Nanopore Technologies [ONT], UK) according to the manufacturer’s instructions. The final libraries were sequenced on the MinION platform (ONT) utilizing an R10.4.1 Flow Cell (FLO-MIN114, ONT).

The nanopore sequencing datasets were subsequently analyzed to determine the methylation profiles, with a specific focus on the *M. FcoHsdM* methylation motif GAAAN8TGG. Following our previous study [54], the bioinformatics analysis included the following steps: (i) raw sequencing data were processed using Dorado for simultaneous basecalling and modification calling; (ii) the resulting modified BAM (modBAM) files were analyzed using Modkit (https://github.com/nanoporetech/modkit) to extract and estimate methylation frequencies at single-base resolution; (iii) the methylation level at each genomic site was calculated using the formula:

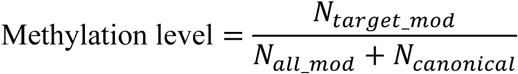

where N_target_mod_ is the number of calls classified as the target modification, N_all_mode_ is the total number of modified calls, and N_canonical_ is the number of unmodified canonical calls; and (iv) the genomic coordinates of all GAAAN_8_TGG motifs were extracted, and the specific methylation levels of adenine residues within these motifs were compared between the WT and Δ*M. FcoHsdM* strains to validate the loss of methylation resulting from the *M. FcoHsdM* gene knockout.

#### Extracellular protein extraction and digestion

Sample collection followed the same procedure as the RNA-seq protocol. Cell cultures were harvested by centrifugation at 5000 rpm, 4 °C for 10 min. The supernatant thus obtained was passed through a membrane filter (pore size, 0.22μm) to remove remaining cells. The filtrate was lyophilized, dissolved in an SDS-free lysis buffer containing 1 × inhibitor cocktail. The mixture was mechanically homogenized and then centrifuged at 25,000 g for 15 min at 4 °C. The proteins in the supernatant were transferred to new centrifuge tubes and 10 mM DTT was added for deoxidation, followed by incubation at 37 °C for 30 min. Thereafter, 55 mM iodoacetamide (IAM) was added, and the mixture was kept in the dark for 45 min. Proteins were then precipitated using five volumes of cold acetone and stored at −20 °C for 2 h, followed by centrifugation at 25,000 g for 15 min at 4 °C, and the supernatant was discarded. The protein pellet was air-dried and treated with SDS-free protein lysate, followed by centrifugation at 25,000 g for 15 min at 4 °C. Protein concentrations were determined by Bicinchoninic acid (BCA) method by BCA Protein Assay Kit (ThermoFisher Scientific, USA).

#### Sample preparation and LC-MS/MS analysis

Proteins (20 μg) were digested with Trypsin (1:50 w/w) at 37 °C overnight. The resulting peptide mixture was separated using a Vanquish Neo UHPLC system (Thermo Fisher Scientific) equipped with an EASY-Spray™ HPLC column (150 μm × 15 cm). The LC gradient was set from 4% to 99% buffer B (80% acetonitrile, 0.1% formic acid) over 13 minutes. Mass spectrometry analysis was performed on an Astral mass spectrometer (Thermo Fisher Scientific) in Data-Independent Acquisition (DIA) mode. The MS1 scan range was 380–980 m/z with a resolution of 240,000 and a maximum injection time (MIT) of 5 ms. The DIA isolation window was set to 300 windows across the MS1 mass range. Fragmentation was performed using higher-energy collisional dissociation (HCD) with a normalized collision energy (NCE) of 25, and the MS2 MIT was set to 3 ms.

#### Protein identification and quantification

Raw DIA mass spectrometry data were processed using the FragPipe computational platform (https://fragpipe.nesvilab.org/). The spectra were searched against the UniProtKB database (uniprotkb_Flavobacterium_columnare_2025_11_17.fasta) using the ultrafast MSFragger search engine [55]. The false discovery rate (FDR) was strictly controlled at < 1% at both the peptide and protein levels to ensure reliable identification. Differential protein expression between groups was evaluated using Welch’s t-test. Proteins meeting the criteria of a fold change > 1.5 and a P-value < 0.05 were defined as significantly differentially expressed. The identified proteins were functionally annotated using the GO and KEGG databases. Differentially expressed proteins (DEPs) were subjected to GO and KEGG pathway enrichment analysis (*P* < 0.05).

#### Statistical analyses

All experiments were conducted with at least three biological replicates. Data were presented as mean ± standard deviation. Statistical analyses were performed using SPSS 16.0. Significance was determined by Student’s t-test or one-way ANOVA followed by LSD test. *P < 0.05* was considered statistically significant. GraphPad Prism version 10 (GraphPad Software, LLC) was also used to compute statistical tests.

---

**Table S1.**
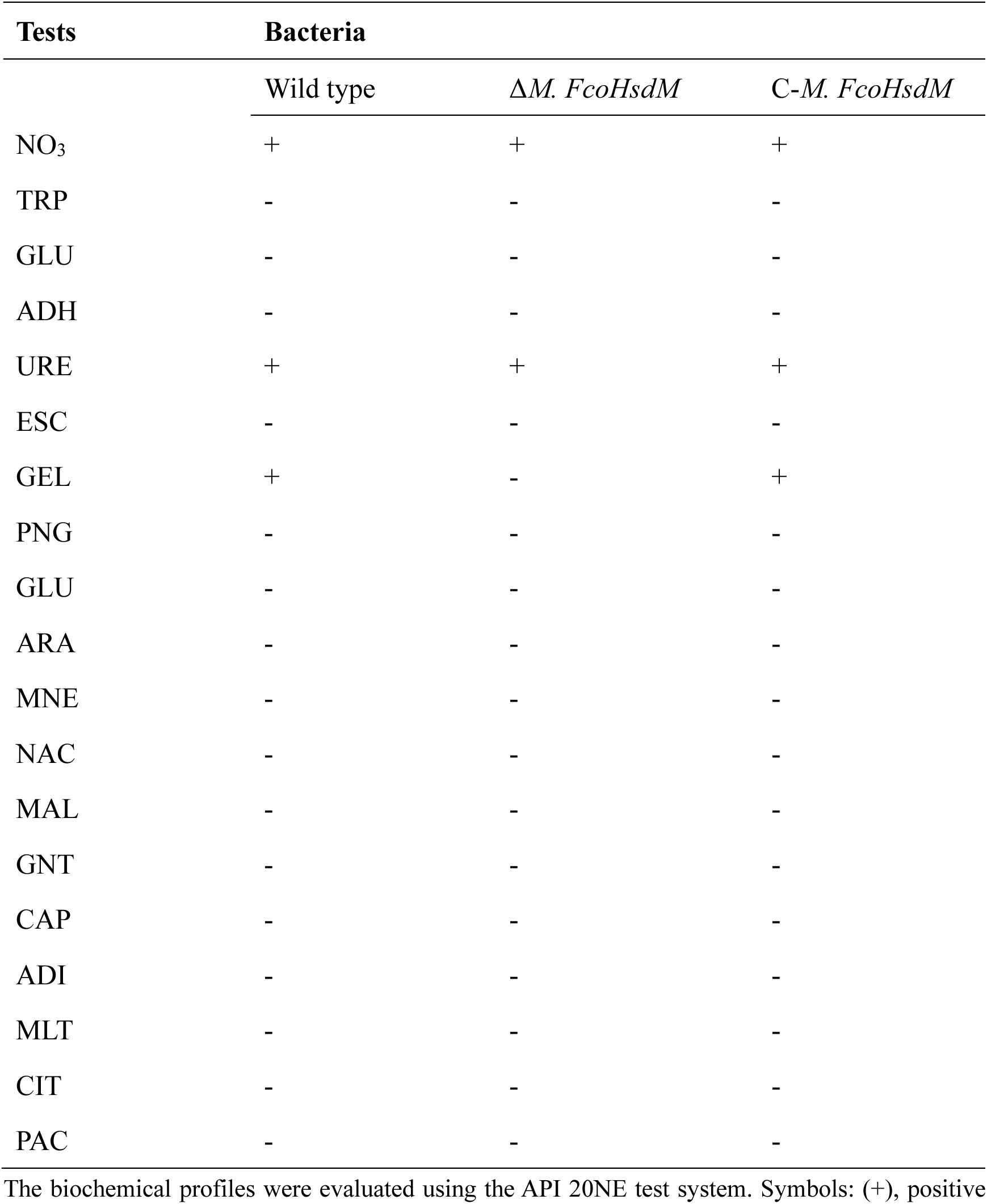
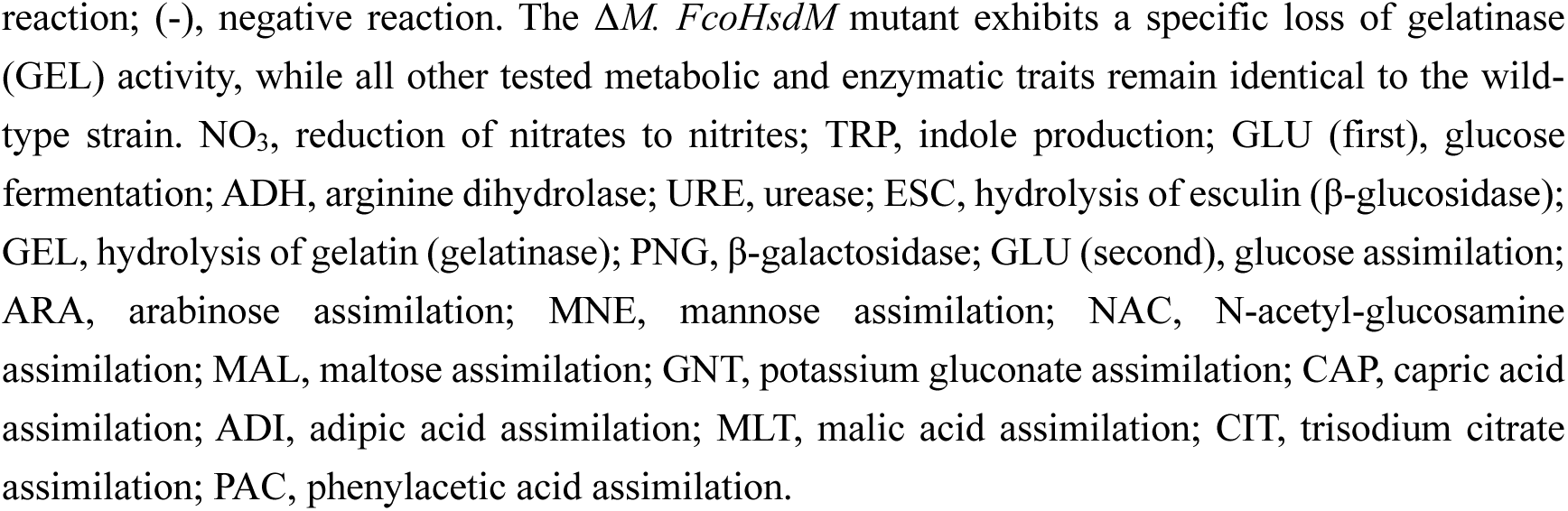
Biochemical characterization of wild-type *F. columnare* and Δ*M. FcoHsdM* strains.

| Tests | Bacteria |  |  |
| --- | --- | --- | --- |
| | Wild type | $\Delta M.$ *FcoHsdM* | *C-M. FcoHsdM* |
| NO<sub>3</sub> | + | + | + |
| TRP | - | - | - |
| GLU | - | - | - |
| ADH | - | - | - |
| URE | + | + | + |
| ESC | - | - | - |
| GEL | + | - | + |
| PNG | - | - | - |
| GLU | - | - | - |
| ARA | - | - | - |
| MNE | - | - | - |
| NAC | - | - | - |
| MAL | - | - | - |
| GNT | - | - | - |
| CAP | - | - | - |
| ADI | - | - | - |
| MLT | - | - | - |
| CIT | - | - | - |
| PAC | - | - | - |

The biochemical profiles were evaluated using the API 20NE test system. Symbols: (+), positive
reaction; (-), negative reaction. The $\Delta M. FcoHsdM$ mutant exhibits a specific loss of gelatinase (GEL) activity, while all other tested metabolic and enzymatic traits remain identical to the wild-type strain. NO<sub>3</sub>, reduction of nitrates to nitrites; TRP, indole production; GLU (first), glucose fermentation; ADH, arginine dihydrolase; URE, urease; ESC, hydrolysis of esculin ( $\beta$ -glucosidase); GEL, hydrolysis of gelatin (gelatinase); PNG, $\beta$ -galactosidase; GLU (second), glucose assimilation; ARA, arabinose assimilation; MNE, mannose assimilation; NAC, N-acetyl-glucosamine assimilation; MAL, maltose assimilation; GNT, potassium gluconate assimilation; CAP, capric acid assimilation; ADI, adipic acid assimilation; MLT, malic acid assimilation; CIT, trisodium citrate assimilation; PAC, phenylacetic acid assimilation.

**Table S2.** Antibiotic sensitivity of wild-type and Δ*M. FcoHsdM*.

| Antibiotic | Resistant<br>mm | Intermediate<br>mm | Susceptible<br>mm | WT | $\Delta M.$<br>*FcoHsdM* |
| --- | --- | --- | --- | --- | --- |
| Penicillin | <19 | 20-27 | >27 | S | S |
| Oxacillin | $\leq 10$ | 11-12 | $\geq 13$ | R | R |
| Madumycin | $\leq 13$ | 14-17 | $\geq 18$ | S | S |
| Carbenicillin | <19 | 20-22 | >23 | S | S |
| Piperacillin | <17 | 18-20 | >21 | S | S |
| Cefalexin | $\leq 14$ | 15-17 | $\geq 18$ | S | S |
| Ofloxacin | $\leq 12$ | 13-15 | $\geq 16$ | S | S |
| Cefuroxime | $\leq 14$ | 15-17 | $\geq 18$ | S | S |
| Florfenicol | $\leq 12$ | 13-17 | $\geq 18$ | S | S |
| Ceftriaxone | $\leq 13$ | 14-21 | $\geq 22$ | S | S |
| Amikacin | $\leq 14$ | 15-16 | $\geq 17$ | S | S |
| Vancomycin | $\leq 9$ | 10-11 | $\geq 12$ | S | I |
| Kanamycin | $\leq 13$ | 14-17 | $\geq 18$ | S | S |
| Neomycin | $\leq 12$ | 13-16 | $\geq 17$ | S | S |
| Furazolidone | $\leq 14$ | 15-16 | $\geq 17$ | S | S |
| Chloramphenicol | $\leq 12$ | 13-17 | $\geq 18$ | S | S |
| Minocycline | $\leq 14$ | 15-18 | $\geq 19$ | S | S |
| Clindamycin | $\leq 14$ | 15-20 | $\geq 21$ | S | S |
| Cefoperazone | $\leq 15$ | 16-20 | $\geq 21$ | S | S |
| Ceftazidime | ≤14 | 15-17 | ≥18 | S | S |
| Cefradine | ≤14 | 15-17 | ≥18 | S | S |

**Table S3.**
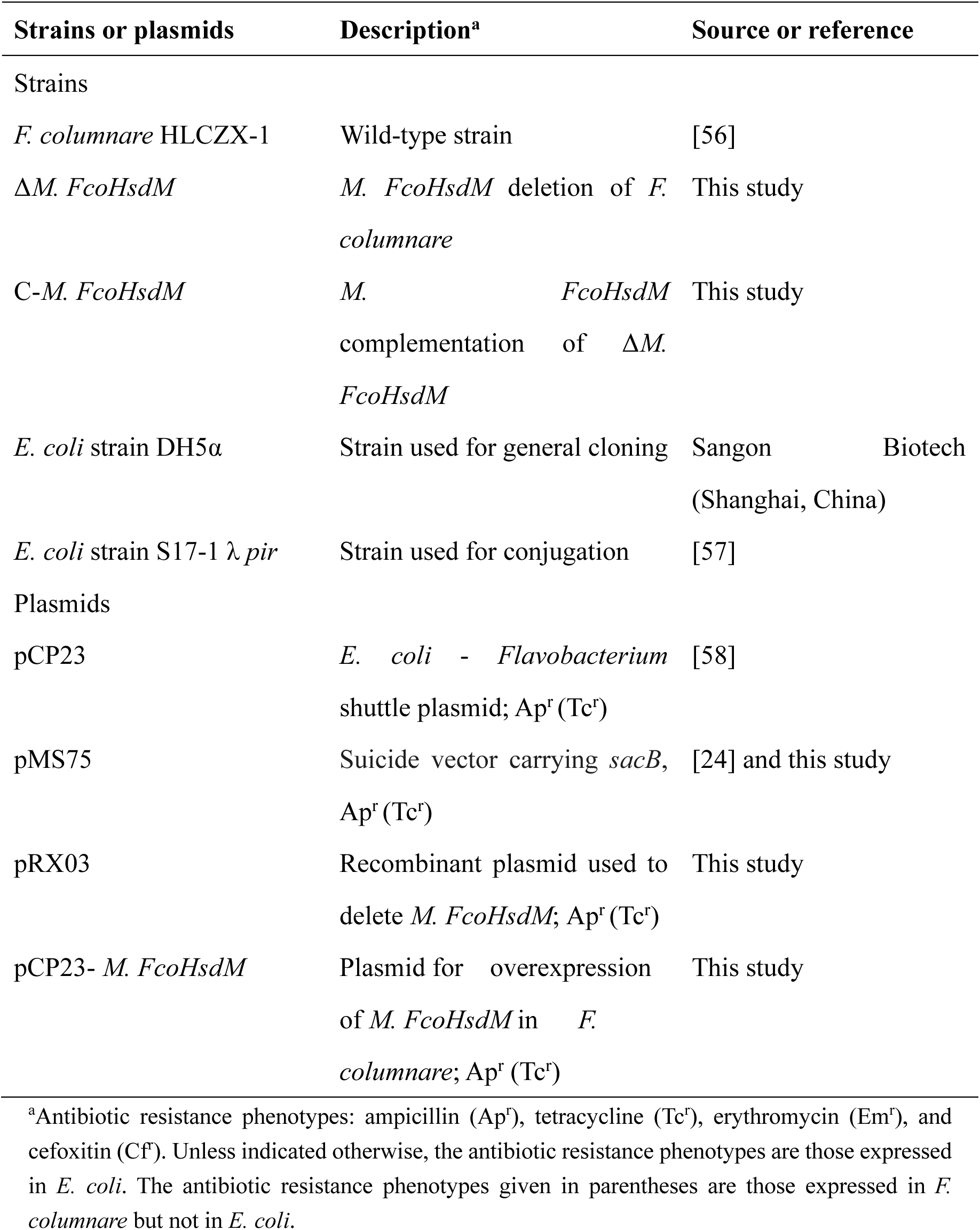
Bacterial strains and plasmids used in this study.

| Strains or plasmids | Description<sup>a</sup> | Source or reference |
| --- | --- | --- |
| Strains |  |  |
| *F. columnare* HLCZX-1 | Wild-type strain | [56] |
| $\Delta M. FcoHsdM$ | *M. FcoHsdM* deletion of *F. columnare* | This study |
| C-*M. FcoHsdM* | *M. FcoHsdM* complementation of $\Delta M. FcoHsdM$ | This study |
| *E. coli* strain DH5α | Strain used for general cloning | Sangon Biotech (Shanghai, China) |
| *E. coli* strain S17-1 $\lambda$ *pir* | Strain used for conjugation | [57] |
| Plasmids |  |  |
| pCP23 | *E. coli* - *Flavobacterium* shuttle plasmid; Ap<sup>r</sup> (Tc<sup>r</sup>) | [58] |
| pMS75 | Suicide vector carrying *sacB*, Ap<sup>r</sup> (Tc<sup>r</sup>) | [24] and this study |
| pRX03 | Recombinant plasmid used to delete *M. FcoHsdM*; Ap<sup>r</sup> (Tc<sup>r</sup>) | This study |
| pCP23- *M. FcoHsdM* | Plasmid for overexpression of *M. FcoHsdM* in *F. columnare*; Ap<sup>r</sup> (Tc<sup>r</sup>) | This study |

<sup>a</sup>Antibiotic resistance phenotypes: ampicillin (Ap<sup>r</sup>), tetracycline (Tc<sup>r</sup>), erythromycin (Em<sup>r</sup>), and cefoxitin (Cf<sup>r</sup>). Unless indicated otherwise, the antibiotic resistance phenotypes are those expressed in *E. coli*. The antibiotic resistance phenotypes given in parentheses are those expressed in *F. columnare* but not in *E. coli*.

**Table S4.**
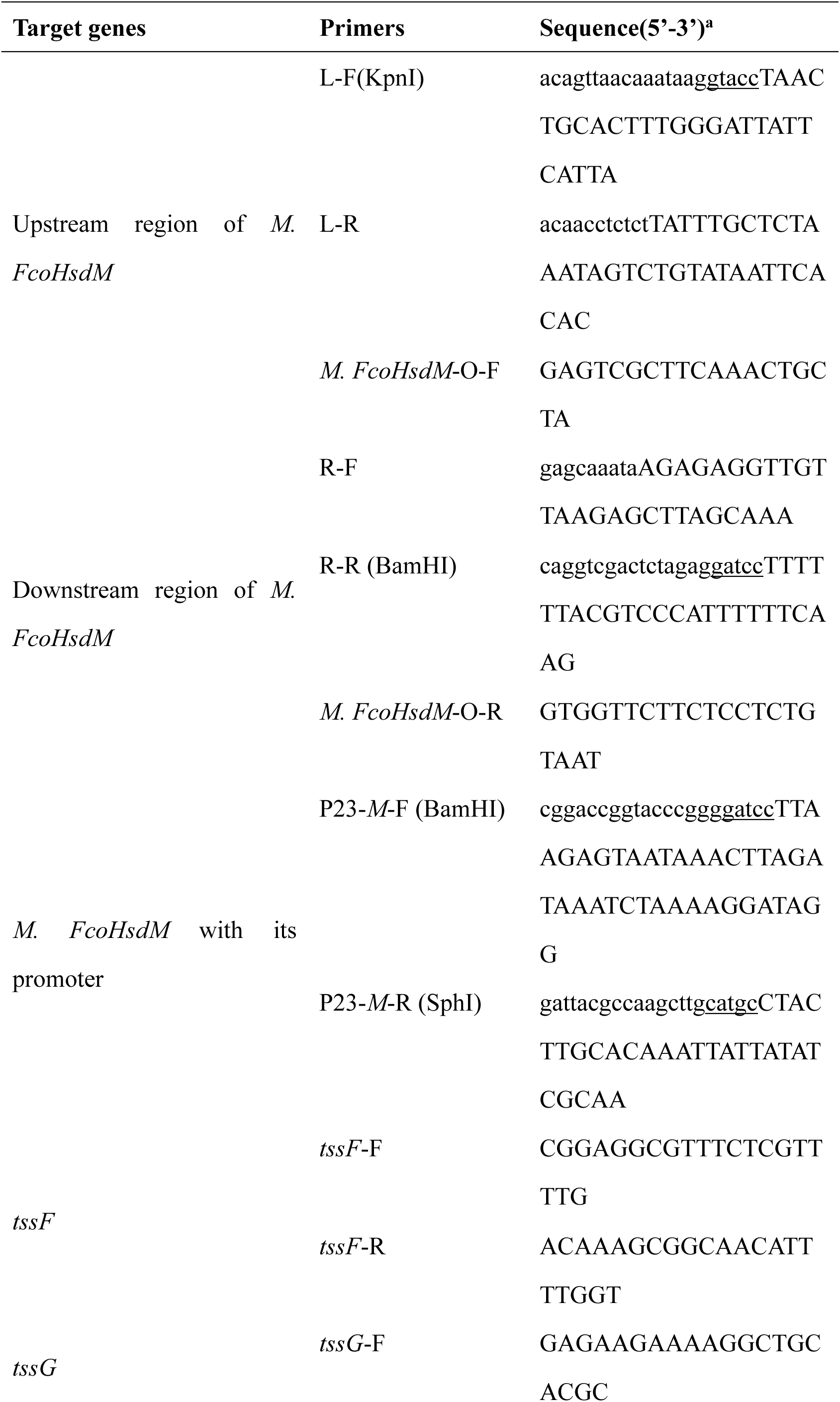
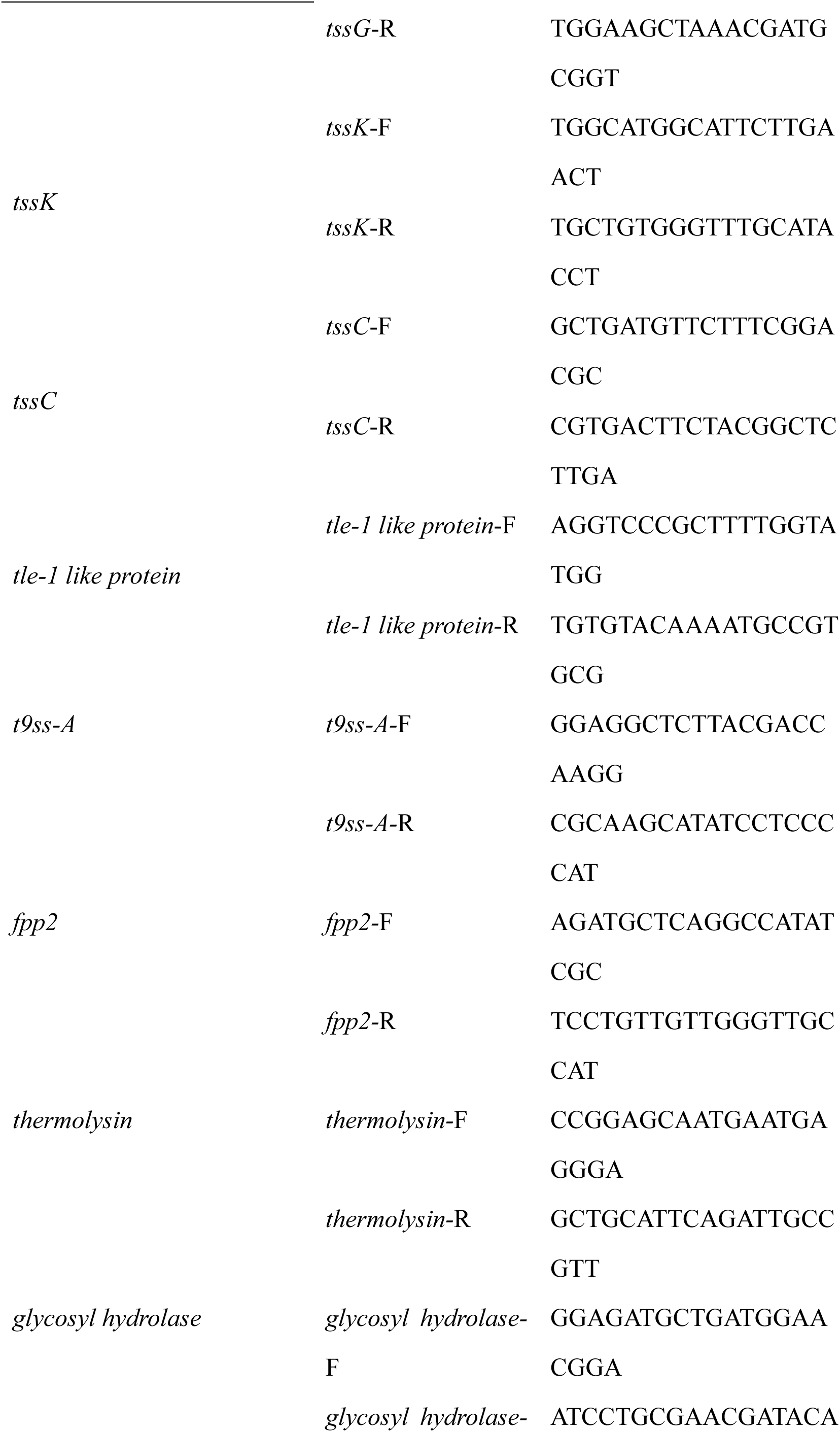
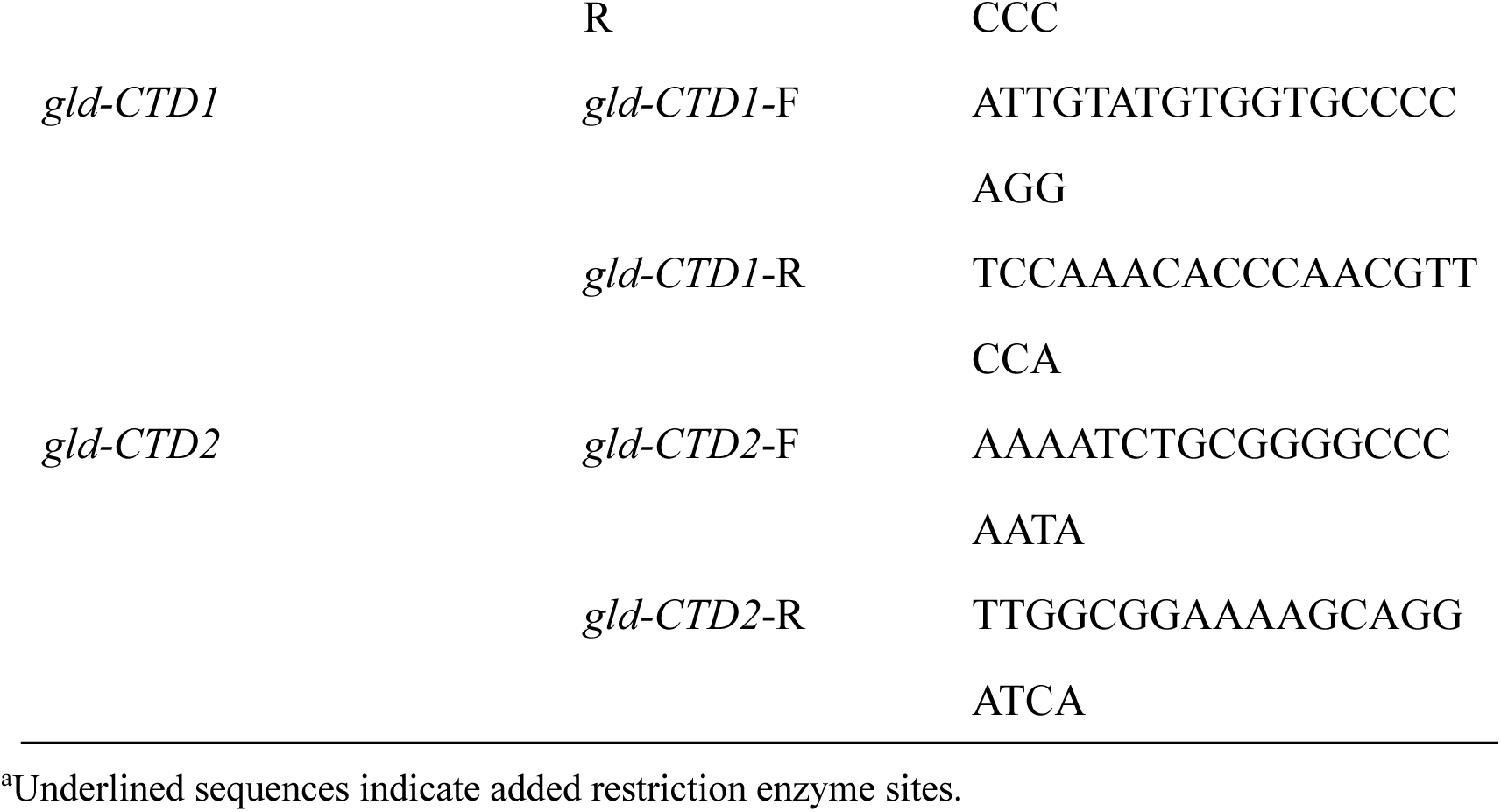
Sequences of primers used in this research.

## Supplementary figures

**Figure S1.**
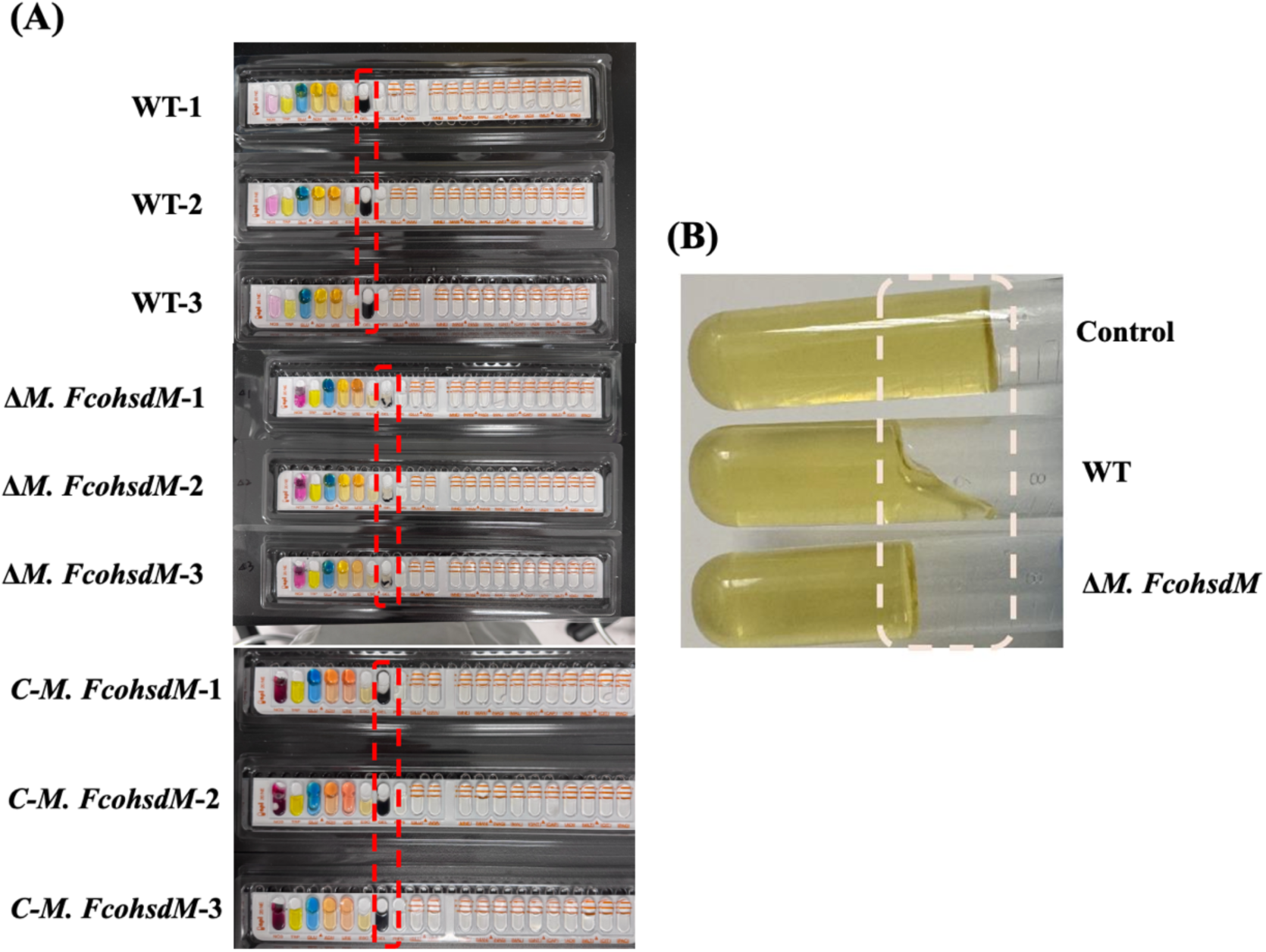
Loss of gelatinase activity in the Δ*M. FcoHsdM* mutant. **(A)** Biochemical characterization using the API 20NE test system. The red dashed boxes highlight the gelatinase (GEL) reaction wells. The wild-type (WT) strains (three biological replicates) exhibit a positive reaction (black pigmentation), whereas the Δ*M. FcoHsdM* mutant strains show a negative reaction. **(B)** Nutrient gelatin stab assay confirming the impaired proteolytic capacity. The WT strain completely liquefied the gelatin medium, indicating active gelatinase secretion, while the Δ*M. FcoHsdM* mutant and the uninoculated control failed to cause liquefaction (the medium remained solid).

**Figure S2.**
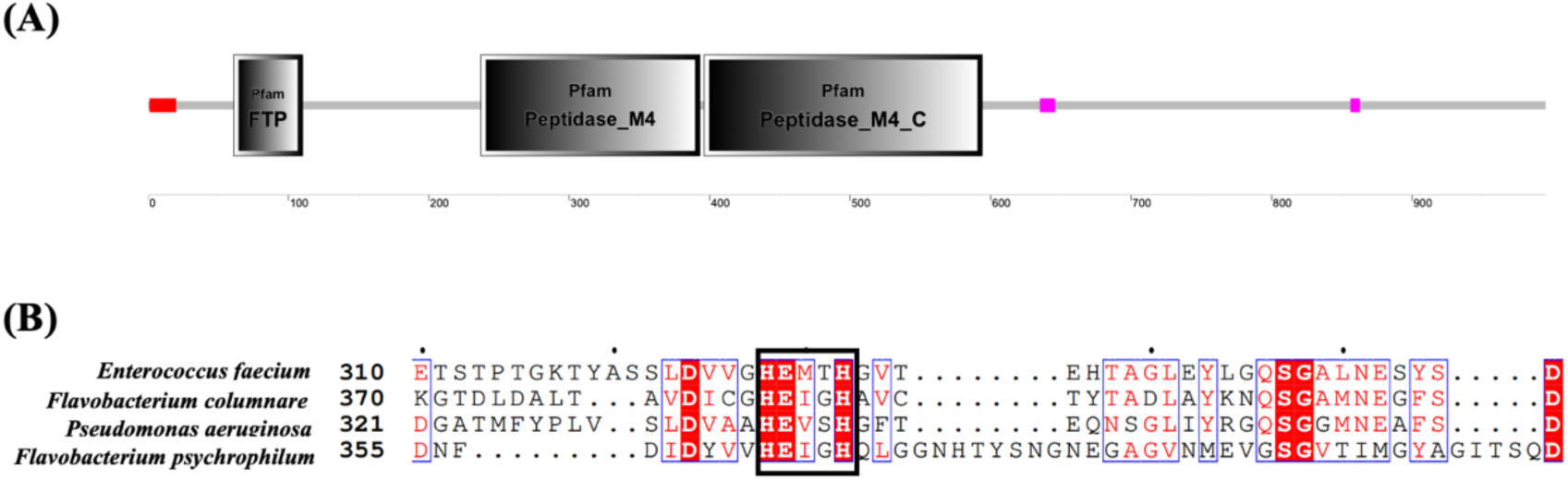
Domain architecture and sequence alignment of the putative gelatinase Fco1. **(A)** Predicted structural domains of Fco1 from *F. columnare*, showing the N-terminal FTP domain, the central Peptidase_M4 catalytic domain, and the C-terminal Peptidase_M4_C domain. **(B)** Multiple sequence alignment of the Fco1 catalytic region with homologous metalloproteases from other bacterial species. Identical and highly conserved amino acid residues are highlighted with a red background and red font, respectively. The solid black box indicates the conserved zinc-binding motif (HEXXH) characteristic of M4 family metalloproteases.

**Figure S3.**
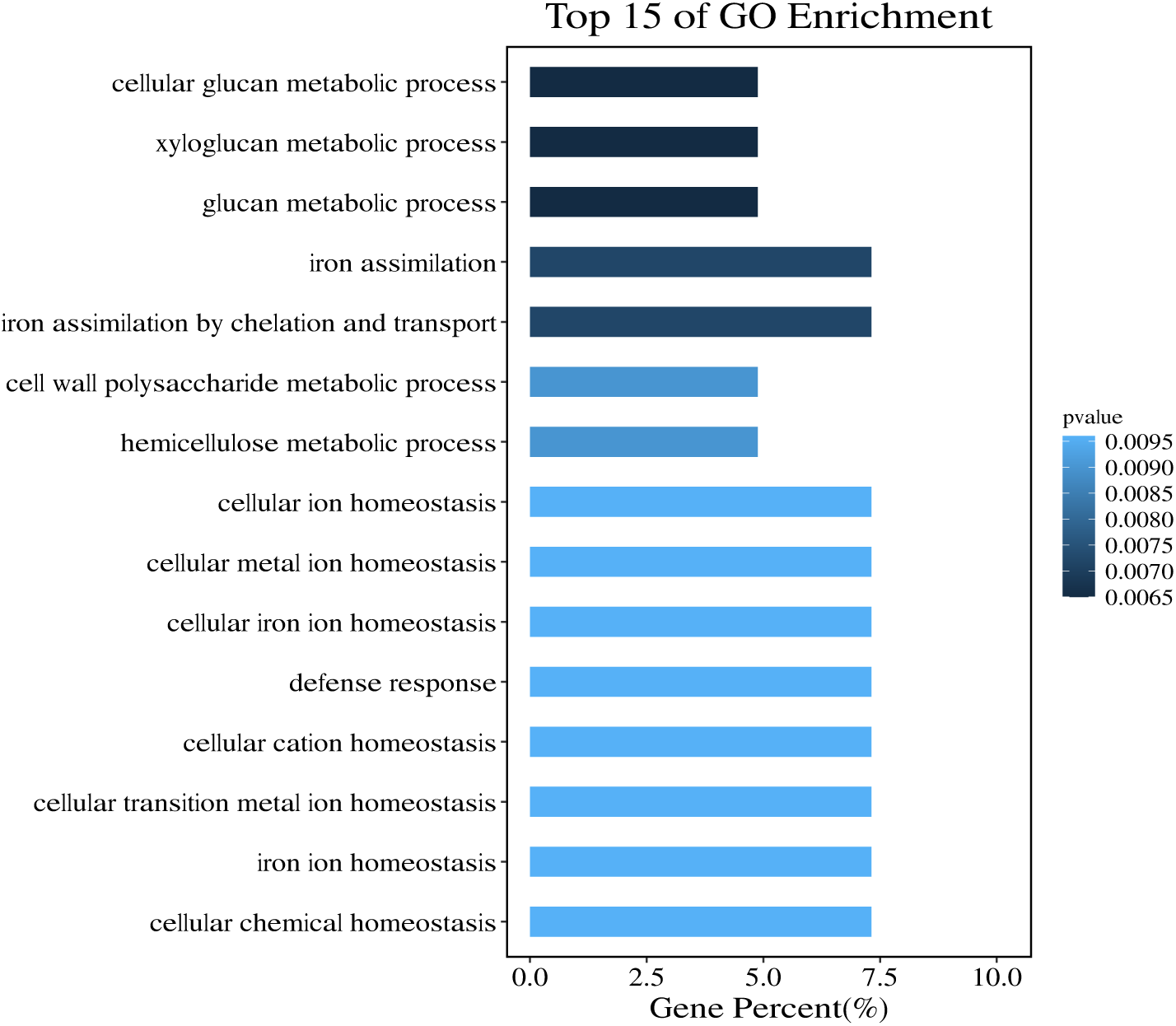
GO enrichment analysis of the differentially expressed genes (DEGs) containing the M. FcoHsdM methylation motif.

